# Microbiota–modulated enteric neuron translational profiling uncovers a CART+ glucoregulatory subset

**DOI:** 10.1101/2020.03.09.983841

**Authors:** Paul A Muller, Marc Schneeberger, Fanny Matheis, Zachary Kerner, Daniel Mucida

## Abstract

Microbial density and diversity increase towards the distal intestine, affecting tissue physiology, metabolism, and function of both immune and nervous systems. Intrinsic enteric–associated neurons (iEAN) continuously monitor and modulate intestinal functions, including nutrient absorption and motility. Through molecular, anatomic and functional approaches, we characterized the influence of the microbiota on iEAN. We found that iEAN are functionally adapted to the intestinal segment they occupy, with a stronger microbiota influence on distal intestine neurons. Chemogenetic characterization of microbiota-influenced iEAN identified a subset of viscerofugal CART+ neurons, enriched in the distal intestine, able to modulate feeding through insulin-glucose levels. Retro- and anterograde tracing revealed that CART+ viscerofugal neurons send axons to the gut sympathetic ganglion and are synaptically connected to the liver and pancreas. Our results demonstrate a region-specific adaptation of enteric neurons and indicate that specific iEAN subsets are capable of regulating host physiology independently from the central nervous system.

**One Sentence Summary:** Microbes impact regionally defined intrinsic enteric neuron translatomes, including a novel CART+ glucoregulatory viscerofugal population.

## Main Text

EAN comprise a numerous and heterogeneous population of neurons within the gastrointestinal (GI) tract that monitor and respond to various environmental cues such as mechanical stretch and luminal metabolites (*1, 2*). The vast majority of luminal stimuli are derived from the diet and commensal microbes, which may be sensed directly by EAN fibers positioned along the intestinal epithelium. Luminal perturbations can also be transmitted to EAN indirectly, via signals derived from epithelial, glial, or immune cells inhabiting the same compartment (*1, 3*). Intrinsic EAN (iEAN), which comprise a component of the enteric nervous system (ENS), are neural crest–derived and organized in two distinct layers, the myenteric or Auerbach’s plexus and submucosal or Meissner’s plexus (*2*). iEAN can operate autonomously and are primarily tasked with modulation of intestinal motility and secretory function (*2*). Recent studies have demonstrated that the gut microbiota influence the basal activity of intestine–associated cells, including the excitability of EAN and the activation state of immune cells (*2-5*). Additionally, microbial dysbiosis has a potential role in a host of metabolic disorders including obesity and diabetes (*6, 7*). Yet, whether the metabolic effects of the microbiota are mediated through the nervous system is still not known. These studies highlight the impact of the gut microbiota on EAN and key mammalian physiological processes, however the cellular circuits and molecular components that mediate gut-EAN or gut–brain communication remain poorly understood. We sought to determine how the microbiota impacts iEAN to better characterize their role in host physiology.

To profile iEAN, we opted for a translating ribosomal affinity purification (TRAP) approach(*8*); cell type– specific mRNA profiling that gives information on what is being actively translated within the cell and bypasses the need for tissue fixation or single–cell suspension, avoiding possible confounding effects of neuronal dissociation on gene expression. We interbred pan–neuronal *Snap25*^Cre^ with *Rpl22*^lsl-HA^ (RiboTag) mice(*9*), which express a hemagglutinin (HA)–tagged ribosomal subunit 22, allowing immunoprecipitation of actively–translated mRNA. Expression of HA–tagged ribosomes was observed in neurons in the myenteric plexus of the duodenum, ileum, and colon of *Snap25*^RiboTag^ mice (Fig. 1A). We confirmed successful immunoprecipitation (IP) of intact mRNA bound to HA-tagged ribosomes from myenteric neurons in the intestine muscularis. RNA sequencing of bound transcripts revealed significant enrichment of neuronal-specific genes and pathways in *Cre*^+^ animals when compared to *Cre*^−^ controls (fig. S1A-C). The TRAP system allowed the identification of novel enteric neuron markers such as CD9, which was confirmed to be highly expressed in EAN cell bodies and fibers in the myenteric plexus, but not present in other cell types such as enteric glia (fig. S1D, E). TRAP RNA-seq (TRAP-seq) analysis of iEAN and extrinsic EAN (nodose, NG; celiac–superior mesenteric, CG-SMG; and dorsal root ganglion, DRG)(*10*) suggested that iEAN possess a distinct translational profile (Fig. 1B, C). iEAN expressed more neuropeptide transcripts compared to sensory and sympathetic ganglia, which had increased expression of genes involved in nociception and catecholamine production, respectively (Fig. 1D, fig. S2A-C).

**Fig. 1.**
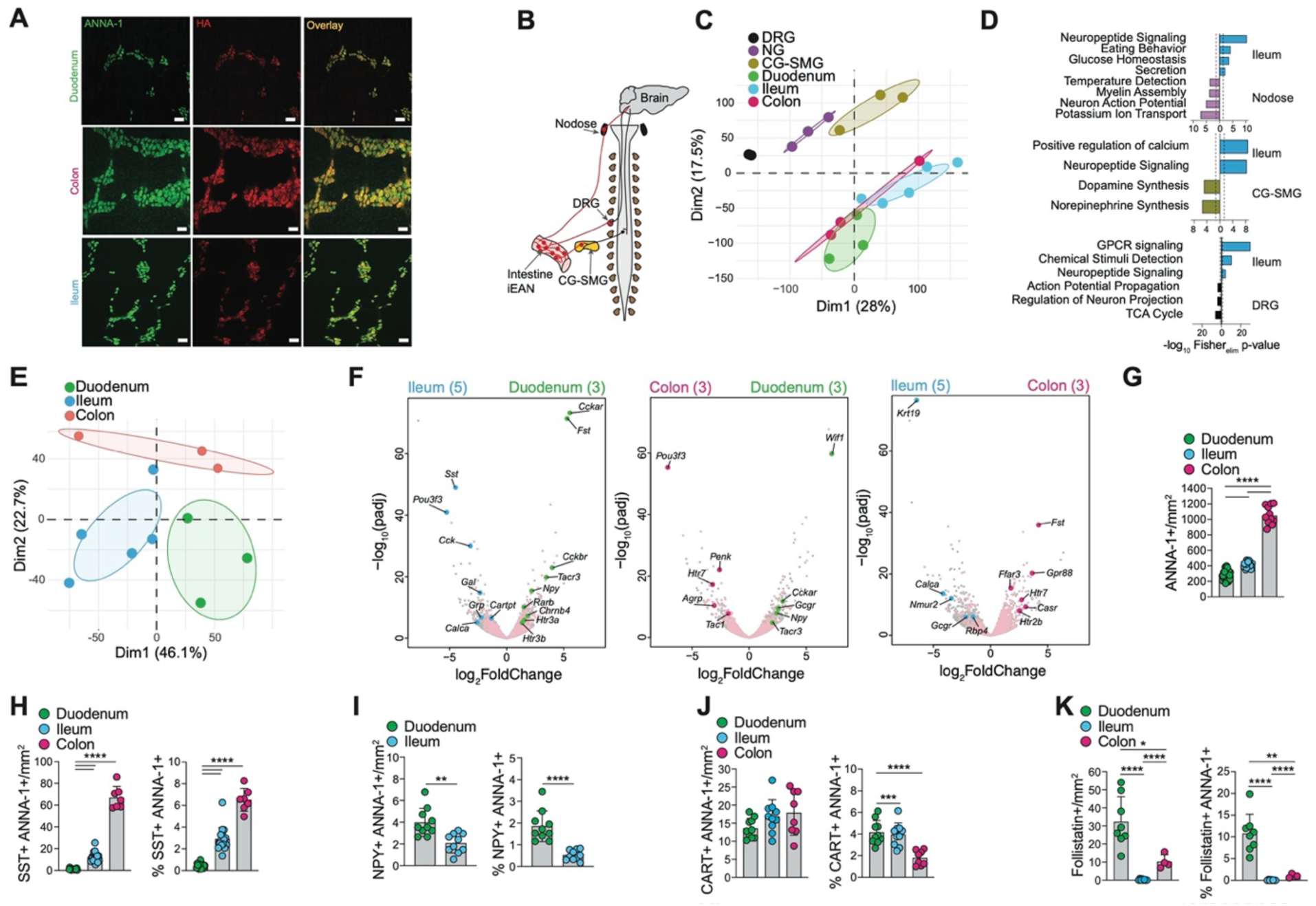
TRAP-seq profiling of iEAN reveals anatomical imprinting. (**A**) Representative whole-mount immunofluorescence (IF) image of myenteric plexus neurons from the duodenum, ileum, and colon of *Snap25*^RPL22-HA^ SPF mice using anti-hemagglutinin (HA) and anti-neuronal nuclear (ANNA-1) antibodies. Scale bars = 50 μm. Images representative of at least n=3. (**B**) Anatomical diagram of intrinsic and extrinsic EAN. Red lines highlight relevant ganglia and fibers. DRG = dorsal root ganglion, CG-SMG = celiac-superior mesenteric ganglion. (**C**) Principal Component Analysis of translatomes from nodose ganglion, celiac-superior mesenteric ganglion (CG-SMG), dorsal root ganglion (DRG), duodenum, ileum, and colon EAN of C57BL/6J SPF mice. (**D**) Gene ontology pathways from TopGO analysis of differentially-expressed genes (log2 Fold Change > 1, padj < 0.05), enriched in the ileum (blue) vs nodose ganglion (purple), CG-SMG (gold), and DRG (black). Dashed lines represent threshold of significance (1.3), calculated by Fisher’s test with an elimination algorithm. (**E**) Principal Component Analysis of translatomes from the duodenum, ileum, and colon EAN of C57BL/6J SPF mice. Immunoprecipitated (IP) transcripts from myenteric iEAN (log2 Fold Change > 1, padj < 0.05) were used to generate the list of genes for comparison between all groups. (**F**) Volcano plots of differentially-expressed genes between myenteric iEAN populations in the duodenum, ileum and colon. Grey dots highlight all genes analyzed by differential expression analysis. Pink dots highlight all IP-enriched genes from each pair of iEAN. Green, blue, and magenta dots represent differentially expressed genes between duodenum, ileum, and colon, respectively. Sample numbers are indicated in parentheses. (**G**) Number of myenteric plexus iEAN per mm^2^ in the duodenum, ileum, and colon of C57BL/6J SPF mice. **** *P* < 0.0001 as calculated by unpaired t-test. (**H-K**) Number and percentage of somatostatin (SST)+ (H), neuropeptide Y (NPY)+ (I), CART+ (J), and follistatin (FST)+ (K) myenteric plexus iEAN per mm^2^ in the duodenum, ileum, and colon of C57BL/6J SPF mice. * *P* < 0.05, ** *P* < 0.01, *** *P* < 0.001, **** *P* < 0.0001 as calculated by unpaired t-test.

Comparison between translational profiles of myenteric neurons isolated from the duodenum, ileum, and colon indicated that iEANs segregate based on their anatomical location (Fig. 1E, fig. S3A-C). Compartmentalized translational profiles of myenteric neurons are consistent with the anatomically distinct functions of the corresponding segments of the intestine. The proximal small intestine is highly absorptive, enriched with enteroendocrine cell (EEC) subsets associated with lipid and nutrient detection (*11*). We found that duodenal iEAN, in comparison those of the ileum and colon, express significantly higher levels of transcripts encoding receptors involved in the response to proximal EEC–derived signals such as *Cckar, Gcgr*, and *Tacr3*, likely reflecting the duodenum’s predominant role in nutrient absorption (Fig. 1F, fig. S3A-C). The terminal ileum and colon iEAN, in contrast, are enriched in neuropeptide transcripts, such as *Sst, Cartpt, Penk, Grp*, and *Tac1*, which are thought to be involved in the control of secretomotor processes in the gut (Fig. 1F, fig. S3A-C). We also found enrichment of the transcription factor *Pou3f3* in the ileum and colon, suggesting a unique gene program in the distal intestine (Fig. 1F). Immunofluorescence (IF) analysis confirmed a region-specific compartmentalization at the protein level, reflecting regional differences in neuronal number along the intestine (Fig. 1G, fig S3D). For instance, the neuropeptide somatostatin (SST), involved in the regulation of several GI hormones and smooth muscle contraction (*12*), is highly expressed in the ileum and colon but scarcely expressed by duodenum EAN (Fig. 1H); neuropeptide Y (NPY), typically involved in the regulation of food intake (*12*), was enriched in duodenum EAN (Fig. 1I, fig. S3E, F). We also observed increased numbers of cocaine and amphetamine related transcript (CART) neurons, important for metabolic regulation (*13, 14*), in the distal intestine (Fig 1J, fig. S3G). Finally, we found that the duodenum is particularly enriched in transcripts such as *Fst1* (encoding follistatin 1) and *Wif1* (encoding WNT inhibitory factor 1) as compared to the ileum and colon, suggesting that these neurons may play a role in the regulation of cell proliferation within this region of the intestine. Immunofluorescence analysis of follistatin confirmed prominent FST1+ neurons and nerve fibers in the duodenum that were absent in the ileum and sparse within the colon (Fig. 1K, fig. S3H). These data reveal that the environment of different intestinal regions program a distinct gene profile on iEAN.

Because the density and diversity of the gut microbiota increases from the proximal to distal intestine, we examined whether regionally distinct iEAN translational programs are partially influenced by the microbiota. To address the influence of microbial stimuli on EAN, we first performed AdipoClear (*15*) on whole–mount intestinal tissue followed by light-sheet microscopy to visualize the three-dimensional structure of EAN in the ileum and colon of germ–free (GF) or specific–pathogen free (SPF) mice. The overall organization of iEAN into the two distinct plexuses appeared unaltered between GF and SPF (Fig. 2A, Supplementary Video 1-4). We observed vast mucosal innervation in the small and large intestines of both GF and SPF mice reaching into individual villi with fibers adjacent to the epithelium (Fig. 2A). Analysis of the ileum indicated significant remodeling of nerve fibers reflecting the thin, blunted villi of GF animals, while colonic innervation did not show gross alterations between mice kept under GF and SPF conditions (Fig. 2A, Supplementary Video 1-4). However, quantification of iEAN in the myenteric plexus revealed a significant reduction in the duodenum and ileum of GF mice, while in the colon GF and SPF mice displayed similar numbers (Fig. 2B). To determine whether the microbiota impacts iEAN gene profile along the intestine, we re-derived *Snap25*^RiboTag^ mice under GF conditions (fig. S4A). Analysis of TRAP-seq from duodenum, ileum, and colon muscularis of GF *Snap25*^RiboTag^ mice suggested a significant influence of the microbiota on the compartmentalization of iEAN phenotypes. In SPF mice, principal component analysis segregated proximal and distal intestinal regions. However, ileum, colon and duodenum samples from GF mice all clustered together with the duodenum samples of SPF mice, the region with the lowest microbial density (Fig. 2C). Analysis of the third principal component showed segregation of colon samples from the small intestine, which may reflect the presence of iEAN derived from sacral progenitors in the large intestine (*16*) (fig. S4B). Analysis of GF and SPF datasets using predicting associated transcription factors from annotated affinities (PASTAA) identified CREB amongst the most enriched transcription factors for the colon and ileum in SPF mice (fig. S4C). Because the level of pCREB in neurons is often used as a indirect measure of activation (*17*), we assessed phosphorylation of CREB (pCREB) at serine 133, key to inducing gene transcription, by IF. We found a significant reduction of pCREB in the ileum myenteric plexus of GF compared to SPF mice (fig. S4D,E), demonstrating that iEAN may be hypoexcitable under gnotobiotic conditions, as previously proposed (*18*).

**Fig. 2.**
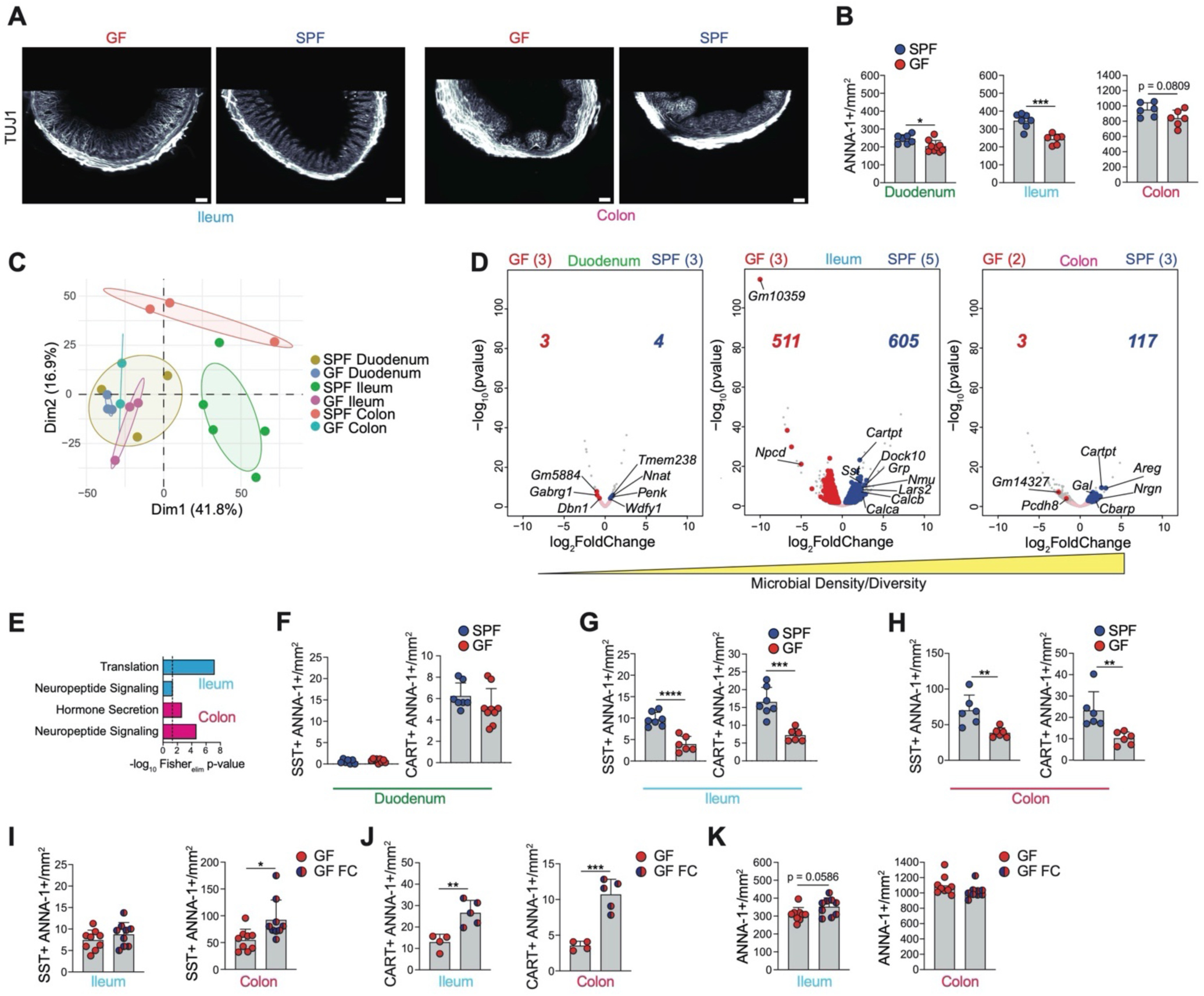
Microbiota impacts iEAN translatome in a compartmentalized manner. (**A**) Light-sheet images of the ileum and colon of AdipoClear-based cleared tissue of C57BL/6J SPF or GF mice stained with anti-β-lll-tubulin (TUJ1) antibodies. Images representative of n=5 for all groups. Scale bar GF ileum = 200 μm, SPF ileum = 200 μm, GF colon = 200 μm, SPF colon = 100 μm. (**B**) Number of myenteric iEAN in the duodenum, ileum, and colon of C57BL6/J GF and SPF mice kept on a GF diet. * *P* < 0.05, *** *P* < 0.001 calculated by unpaired t-test. (**C**) Principal Component Analysis of iEAN from the duodenum, ileum, and colon of C57BL6/J SPF and GF mice. Total comparative gene list generated from IP-enriched transcripts (log2Fold Change > 2.5, padj < 0.05) from each sample. (**D**) Volcano plots of differentially-expressed genes of iEAN from the duodenum, ileum, and colon of C57BL/6J SPF and GF mice. Grey dots highlight all genes analyzed by differential expression analysis. Pink dots highlight all IP-enriched genes from each pair of compaired intestine segments. Red dots and number highlight genes higher in GF samples. Blue dots and number highlight genes significantly higher in SPF samples. Sample numbers are indicated in parentheses.(**E**) Gene ontology pathways, identified by TopGO analysis, of differentially-expressed genes (log2 Fold Change > 1, padj < 0.05), enriched in SPF ileum (blue) and colon (magenta) as compared to respective GF samples. Dashed lines represent threshold of significance (1.3) as calculated by Fisher’s test with an elimination algorithm. (**F-H**) Number of somatostatin (SST)+ and CART+ myenteric iEAN in the duodenum (F), ileum (G), and colon (H) of C57BL/6J GF and SPF mice. ** *P*< 0.01, *** *P* < 0.001, *** *P* < 0.0001 as calculated by unpaired t-test. (**I-K**) Number of somatostatin (SST)+ (I), CART+ (J), and total (K) myenteric iEAN in the ileum (left) and colon (right) of GF mice and GF mice 2 weeks post colonization with microbiota of SPF mice (fecal colonization, GF FC). * *P* < 0.05, ** *P* < 0.01, *** *P*< 0.001 as calculated by unpaired t-test.

Comparison of GF duodenum, ileum, and colon samples also indicated segregation between regions, suggesting that certain features of region-specific iEAN programming are microbiota-independent (fig. S4F). However, in the duodenum, only four genes were significantly upregulated in SPF compared to GF including *Nnat* and *Penk*, involved in neuronal development (*19*) and enkephalin production (*20*), respectively. In the ileum and colon, we detected 605 and 117 differentially expressed genes upregulated, respectively, in SPF as compared to GF groups (Fig. 2D). Among these were genes encoding neuropeptides associated with a neuro-immune crosstalk and with EAN physiological function, such as *Nmu* (*21*), *Sst, Cartpt*, and *Agrp* (colon only) (Fig. 2D, E). SST and CART protein expression changes were confirmed by quantification of immunofluorescence images from SPF and GF mice (Fig. 2F-H, fig. S4G-L). These results establish regional differences as well as the microbial influence on iEAN gene profiles, particularly on their neurochemical coding. To address whether altered neuropeptide levels in GF mice are the result of a developmental defect, we provided adult C57BL/6 GF with age- and sex-matched feces from SPF mice on a matched GF diet (exGF). Colonization of 8-week old GF animals with SPF feces was sufficient to increase the number of SST+ and CART+ neurons in the colon and ileum to levels similar to SPF animals after 2 weeks, as well as a notable increase in the density of SST+ and CART+ nerve fibers (Fig. 2I, J, fig. S4M-P). We also noted that in the ileum there was a trend towards an increase in iEAN numbers, whereas the colon remained unaffected by colonization (Fig 2K), an effect that could be attributed to a developmental defect or the lack of a specific bacteria in the recolonization procedures. Overall, the lack of significant changes in the microbial–sparse duodenum, along with the accumulation of significant changes in iEAN gene expression and neurochemical coding in areas with increased microbial diversity and density, suggest that iEAN regional differences are largely determined by microbiota stimulation.

To define whether microbiota-dependent changes were reversible, or imprinted in iEAN upon initial exposure, we administered antibiotics (vancomycin, ampicillin, metronidazole, and neomycin) in the drinking water of SPF mice for 2 weeks. We detected a significant decrease in the number of iEAN in all three intestinal areas analyzed (Fig. 3A). This neuronal reduction was not permanent, as antibiotic withdrawal for two weeks resulted in the recovery of neuronal numbers to SPF levels, similar to what we observed in GF recolonization experiments (Fig. 3B). We recently described an inflammasome-dependent post–infection neuronal death pathway (*22*). To evaluate whether iEAN loss post microbiota depletion was also dependent on Caspase 11 (Caspase 4 in humans), *Casp1Casp11* (ICE^−/–^) or *Casp11*^−/–^ mice were exposed to Splenda or antibiotic on drinking water. Quantification of iEAN in the ileum of antibiotic–treated mice did not reveal iEAN loss in ICE or *Casp11*^−/–^ mice, suggesting an additional role for Caspase 11 in the maintenance of iEAN during dysbiosis (Fig. 3C, D). Treatment with vancomycin, ampicillin or metronidazole, but not neomycin or single-dose streptomycin, also induced a reduction in total neuronal numbers (Fig. 3E). These results suggest a possible role for specific bacteria in the physiological maintenance of iEAN (Fig. 3E-G). Similar to what we observed in GF mice regarding specific microbiota– modulated neuropeptide pathways, we observed a significant decrease in the number and overall percentage of SST+ and CART+ neurons in the ileum and colon upon antibiotic treatment, while the duodenum remained unchanged (Fig. 3H, I, fig. S5A-E). Short-term microbiota depletion with single-dose streptomycin given to wild-type mice, or continuous broad-spectrum antibiotic-treatment of *Casp1/11*^*-/-*^ and *Casp11*^−/–^ mice failed to significantly impact neuropeptide numbers in the distal intestine (Fig. 3J, K, fig. S5F-J). Together with the results obtained in GF animals, these data establish that specific subsets of iEAN, including iEAN expressing neuropeptides SST and CART, are dependent on the microbiota for their maintenance.

**Fig. 3.**
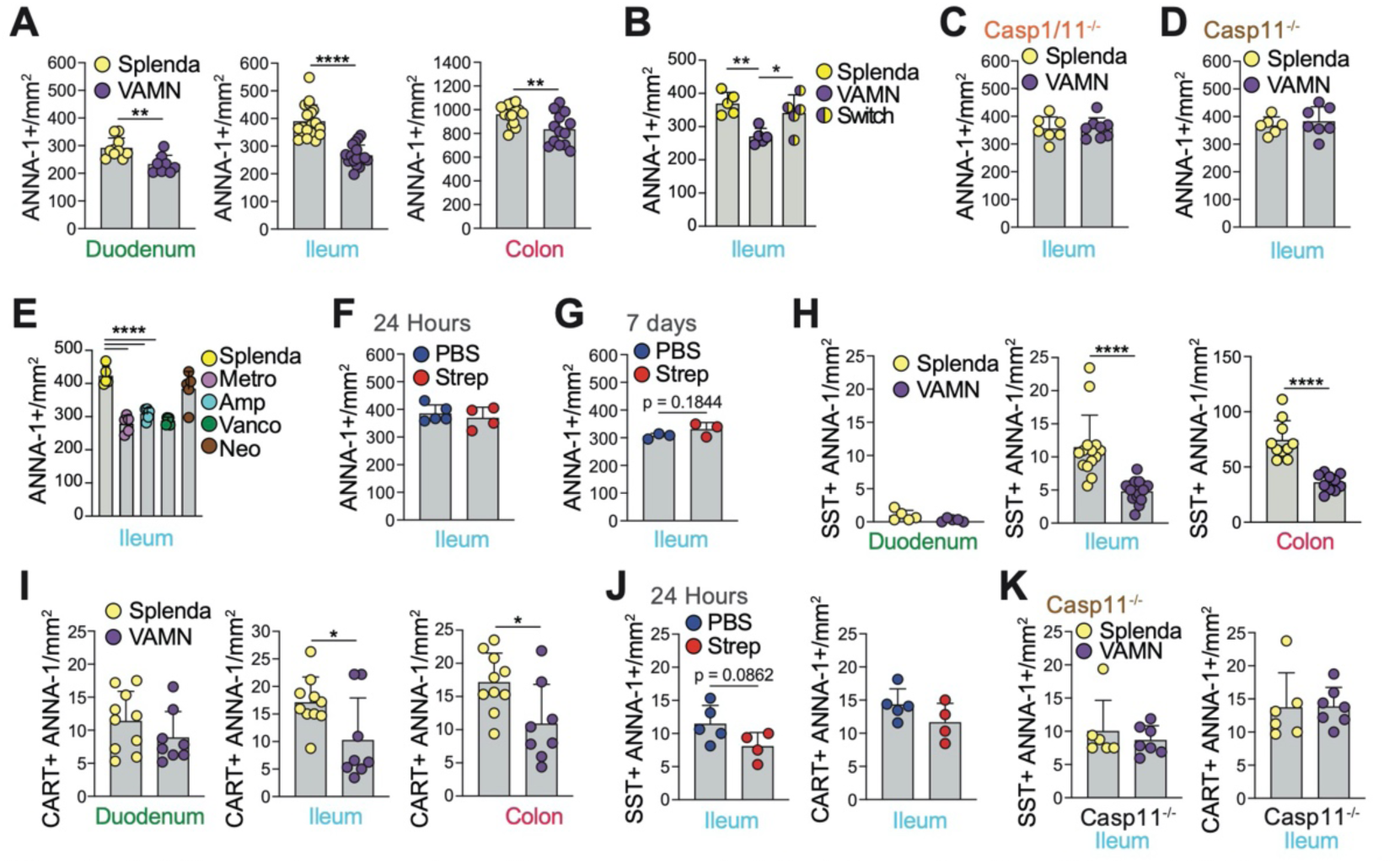
iEAN numbers and profiles are modulated by gut microbiota. (**A**) Number of myenteric iEAN in the duodenum, ileum, and colon of C57BL6/J mice treated with broad-spectrum antibiotics (vancomycin, ampicillin, metronidazole and neomycin - VAMN) administered in Splenda-supplemented drinking water or Splenda (artificial sweetener, control) in the drinking water for two weeks. ** *P* < 0.01, **** *P* < 0.0001 as calculated by unpaired t-test. (**B**) Number of myenteric iEAN in the ileum of C57BL6/J mice treated with either Splenda or VAMN in the drinking water for 4 weeks or VAMN for 2 weeks followed by Splenda for 2 weeks. * *P* < 0.05, ** *P*< 0.01, as calculated by unpaired t-test. (**C, D**) Number of myenteric iEAN in the ileum of *Casp1/11*^*-/-*^ (C) or *Casp11*^*-/-*^ (D) mice treated with either Splenda or VAMN in the drinking water for 2 weeks. (**E**) Number of myenteric iEAN in the ileum of C57BL6/J mice treated with either metronidazole, ampicillin, vancomycin, neomycin, or Splenda in the drinking water for 2 weeks. **** *P* < 0.0001 as calculated by unpaired t-test. (**F**) Number of myenteric iEAN in the ileum of C57BL6/J mice 24 hours post single oral gavage of streptomycin. (**G**) Number of myenteric iEAN in the ileum of C57BL6/J mice 7 days post single oral gavage of streptomycin, p-value calculated by unpaired t-test. (**H**) Number of somatostastin (SST)+ myenteric iEAN in the duodenum, ileum, and colon of C57BL6/J mice treated with broad-spectrum antibiotics (VAMN) or Splenda in the drinking water for two weeks. **** *P* < 0.0001 as calculated by unpaired t-test. (**I**) Numbers of CART+ myenteric iEAN in the duodenum, ileum, and colon of C57BL6/J mice treated with broad-spectrum antibiotics (VAMN) or Splenda in the drinking water for two weeks. * *P* < 0.05 as calculated by unpaired t-test. (**J**) Number of somatostastin (SST)+ and CART+ myenteric iEAN in the ileum of C57BL6/J mice 24 hours post single oral gavage of streptomycin. Non-significant p-value as calculated by unpaired t-test. (**K**) Number of somatostastin (SST)+ and CART+ myenteric iEAN in the ileum of *Casp11*^*-/-*^ mice treated with either VAMN or Splenda in the drinking water for 2 weeks.

We sought to functionally characterize iEAN subsets with specific neuropeptide expression that are modulated by the gut microbiota. To define possible functional outcomes of microbiota–modulated iEANs in GI physiology, we focused on CART+ neurons (enriched in the distal intestine, reversible expression upon microbiota depletion, and unlike SST, not expressed in EECs (*23*)), AGRP+ neurons (enriched in the distal intestine and decreased in GF mice), and on NPY+ neurons (enriched in the duodenum and not affected by the microbiota). These are also three neuropeptides expressed by neuronal populations in the hypothalamus that work in concert to regulate energy balance (*24*), and as such, could potentially play a role in gut-specific circuits influencing feeding behavior. Whole-mount analysis of intestinal muscularis using *in-situ* hybridization confirmed the expression of *Npy* and *Cartpt* in the ileum and colon, and *Agrp* in the mid-colon (fig. S6A-C). We obtained Cre lines corresponding to the three neuropeptides and validated *Cre* expression, along with *Cartpt, Npy*, or *Agrp*, in the periphery using *in situ* hybridization (fig. S6D-G). Because these neuropeptides are known to be expressed both in the periphery (*25-29*) and CNS (*24*), we used a local viral delivery approach to target these neurons and avoid CNS effects. Injection of retrograde adeno-associated virus (AAVrg)-FLEX-tdTomato (*30*) revealed a prominent population of tdTomato+ neurons in the ileum, and colon myenteric plexus of *Cartpt*^Cre^ (*31*) and *Npy*^Cre^ (*32*) mice (Fig. 4A, fig. S6H,I). *Npy*^EAN-tdTomato^ and *Cartpt*^EAN-tdTomato^ neurons displayed considerable innervation of the circular and longitudinal smooth muscle within these segments of the intestine, with *Cartpt*^EAN-tdTomato^ also exhibiting dense inter-ganglionic patterning. Similar to the above *in situ* observations, we found a sparse population of *tdTomato*+ neurons in the mid-colon of *Agrp*^EAN-tdTomato^ mice (*33*), exhibiting muscular and inter-ganglionic innervation (fig. S6J). We were unable to find tdTomato expression in NG, DRG, or CG-SMG in *Cartpt*^EAN-tdTomato^ and *Agrp*^EAN-tdTomato^ mice, while *Npy*^EAN-tdTomato^ mice exhibited a population of tdTomato+ synaptically connected neurons in the CG-SMG (fig. S6K). Of note, we also observed a significant number of tdTomato+ fibers in the CG-SMG of *Cartpt*^Cre^ mice (Fig. 4B, fig. S6L), suggesting targeting of a specific population of viscerofugal neurons, generally defined as mechanosensitive iEAN projecting axons outside of the intestine *(60)*.

**Fig. 4.**
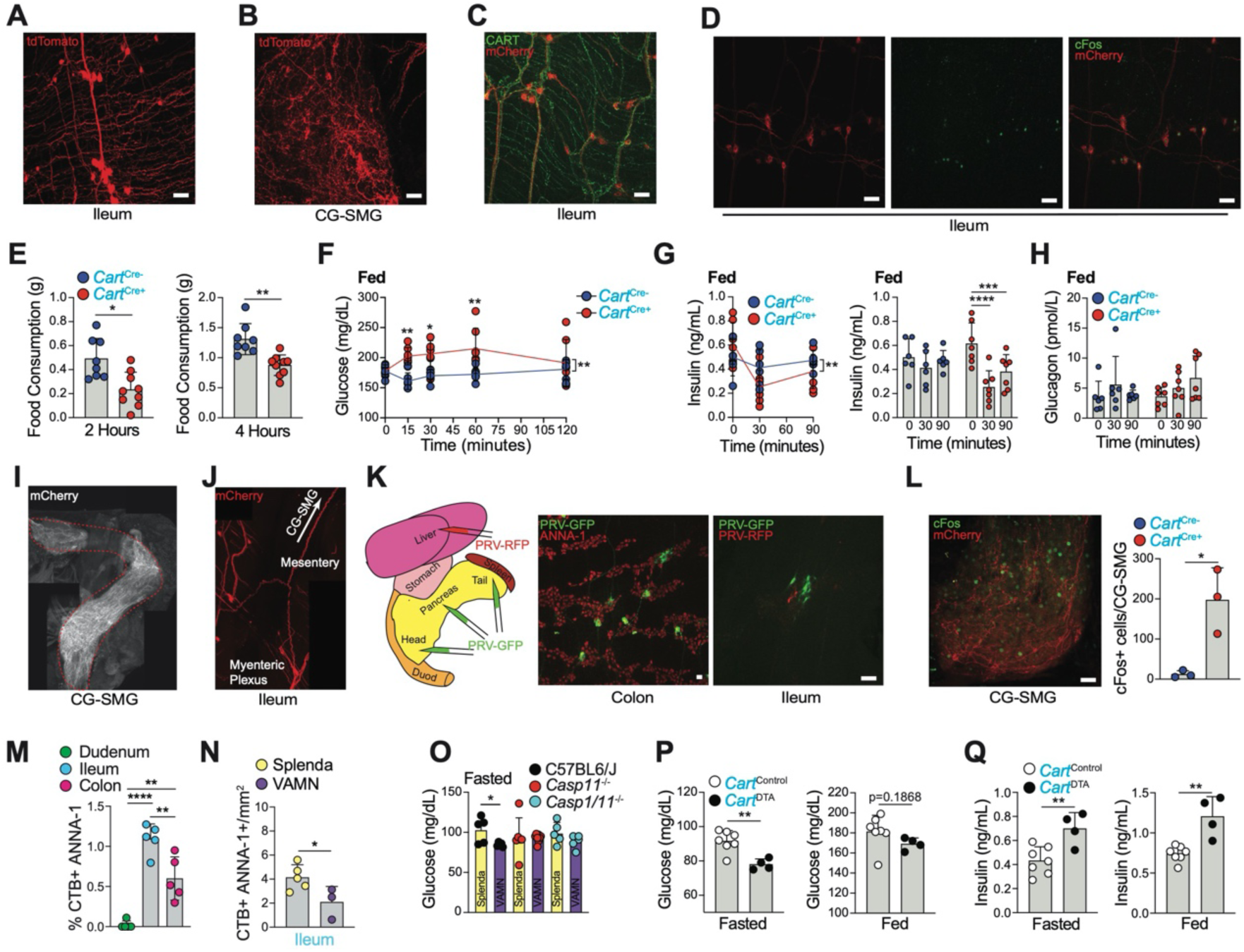
Distal intestine CART+ iEAN are viscerofugal and giucoregulatory. (**A**) Representative whole-mount immunofluorescence (IF) image of the ileum myenteric plexus from *Cart*^Cre^ mice injected with AAVrg-FLEX-tdTomato into the ileum. Scale bars = 50 μm. (**B**) Representative whole-mount IF image of the CG-SMG from *Cart*^Cre^ injected with AAVrg-FLEX-tdTomato into the duodenum, ileum, and colon. Scale bars = 50 μm. (**C**) Reresentative whole-mount IF image of CART+ (green, stained with antibody), and mCherry+ (red, native fluorescence) ileum myenteric iEAN from *Cart*^Cre^ mice injected with AAV9-hSyn-DIO-hM3Dq-mCherry into the ileum. Scale bars = 50 μm. (**D**) Representative whole-mount IF image of cFos+ (green, stained with antibody) and mCherry+ (red, stained with antibody) ileum myenteric iEAN 3 hours post administration of Compound 21 (C21,1mg/kg) to *Cart*^Cre^ mice injected with AAV9-hSyn-DIO-hM3Dq-mCherry into the ileum. Scale bars = 50 μm. (**E-H**) Food consumption at night, 2 hours (left) and 4 hours (right) (E), blood glucose (F), plasma insulin (G), plasma glucagon (H) levels post C21 administration (1mg/kg) to *Cart*^Cre^ mice injected with AAV9-hSyn-DIO-hM3Dq-mCherry into the ileum and colon. * *P*< 0.05, ** *P* < 0.01 as calculated by unpaired t-test (E, G (right) and H), or by two-way ANOVA with Turkey’s multiple comparison’s post hoc test (F, G (left)). *** *P* < 0.001, *** *P* < 0.001. (**I**) Whole-mount immunofluorescence image of the CG-SMG from *Cart*^Cre^ mice injected with AAV9-hSyn-DIO-hM3Dq-mCherry into the ileum and colon. (**J**) Representative whole-mount IF image of the ileum myenteric plexus from *Cart*^Cre^ mice injected with AAV9-hSyn-DIO-hM3Dq-mCherry into the ileum. (**K**) (Left) Scheme of fluorescent PRV injection into the liver and pancreas; (Right) Whole-mount IF image of the colon (left) and ileum (right) myenteric plexus from C57BL6/J mice injected with PRV-GFP into the pancreas and PRV-RFP into the liver. Scale bars = 50 μm. (**L**) (left) cFos (green, stained with antibody) and mCherry (red, native fluorescence) expression in *Cart*^Cre+^ mice expressing AAV5-DIO-hSyn-hM4Di-mCherry in the distal intestine only, 3 hours post-injection of C21 (1mg/kg), (right) Number of cFos+ neurons in the CG-SMG of *Cart*^Cre^ mice injected with AAV9-hSyn-DIO-hM3Dq-mCherry into the ileum and colon 3 hours post injection of C21 (1mg/kg). * *P*< 0.05 as measured by unpaired t-test. Scale bars = 50 μm. (**M** and **N**) Percentage (M) or number (N) of CTB-AF647+ neurons after injection of CTB into the CG-SMG of C57BL6J mice treated with broad-spectrum antibiotics (vancomycin, ampicillin, metronidazole and neomycin - VAMN) administered in Splenda-supplemented drinking water or Splenda (artificial sweetener, control) in the drinking water for two weeks. * *P* < 0.05, ** *P* < 0.01, **** *P* < 0.0001 as calculated by unpaired t-test. (**O**) Blood glucose levels of C57BL6J, *Casp1/11*^−*/*−^ and *Casp11*^−*/*−^ mice treated with broad-spectrum antibiotics (vancomycin, ampicillin, metronidazole and neomycin - VAMN) administered in Splenda-supplemented drinking water or Splenda (artificial sweetener) only in the drinking water for two weeks. * *P* < 0.05 as calculated by unpaired t-test. (**P**) Blood glucose levels of fasted (left) or fed (right) *Cart*^Cre^ mice injected AAV5-mCherry-FLEX-DTA or control virus into the ileum and colon. ** *P* < 0.01 as calculated by unpaired t-test. (**P**) Plasma insulin levels of fasted (left) or fed (right) *Cart*^Cre^ mice injected AAV5-mCherry-FLEX-DTA or control virus into the ileum and colon ** *P* < 0.01 as calculated by unpaired t-test.

To directly assess a potential local GI function played by these three neuropeptide populations, we injected excitatory designer receptor exclusively activated by designer drugs (DREADD) virus (AAV9-FLEX-Syn-hM3Dq-mCherry) into the distal ileum and proximal-mid colon of *Cartpt*^Cre^ and *Npy*^Cre^ mice or into the mid-colon of *Agrp*^Cre^ mice (Fig. 4C, D, fig. S7A). We first performed intestinal motility assays following administration of the DREADD ligand, Compound 21 (C21); we did not observe changes in total intestinal transit in any of the three neuropeptide lines tested (fig. S7B-D). Analysis of feeding behavior also failed to detect robust or consistent changes in both *Npy*^EAN-hM3Dq^ and *Agrp*^EAN-hM3Dq^ mice; however, we observed a significant decrease in food consumption during day feeding at 1 and 2 hours, as well as during night feeding at 2 and 4 hours post C21 in *Cartpt*^EAN-hM3Dq^ mice (Fig. 4E, fig S7E-G). Because CART is expressed by several neuronal populations outside the intestine, including in areas that may influence feeding (*14, 34*), and retrograde transport of AAV9 from the gut has been described (*35*), we examined the NG, DRG, CG-SMG, duodenum and the dorsal motor nucleus of the vagus (DMV) for mCherry+ expression by neuronal populations. We found no clear evidence for hM3Dq expression outside of the distal ileum and proximal colon, indicating that an iEAN-restricted neuronal stimulation can influence feeding (*data not shown*). We evaluated whether the reduction in feeding was related to acute changes in blood glucose or glucoregulatory hormone levels, which can regulate the activity of CNS nuclei controlling feeding behavior (*36-39*). We detected significantly higher blood glucose levels in *Cartpt*^EAN-hM3Dq^ mice injected with C21 as compared to control mice (Fig. 4F, fig. S7H-I). To define whether these changes in basal blood glucose were directly related to typical glucoregulatory mechanisms, we measured insulin and glucagon levels following C21 administration. We found a significant decrease in insulin levels at 30 and 90 minutes post-C21 administration to *Cartpt*^EAN-hM3Dq^ mice, while glucagon levels were only marginally (non-significantly) increased at 90 minutes (Fig. 4G, H). These data indicate that stimulation of distal intestine CART+ neurons result in decrease insulin levels with a subsequent decrease in feeding in mice.

We next asked how CART+ neurons can exert their glucoregulatory function. Imaging analyses confirmed that at least some CART+ neurons in the distal intestine, in particular the ileum, are viscerofugal. These CART+ neurons send axonal projections to the CG-SMG (Fig. 4I, J, Supplementary Video 5), which in turn provides sympathetic innervation to a number of visceral organs, including the pancreas and liver (*40, 41*). Sympathetic innervation of the pancreatic islets can stimulate the release of glucagon and inhibit insulin through adrenergic receptor engagement on alpha and beta cells, respectively (*40, 42*), while sympathetic stimulation of the liver can drive gluconeogenesis (*41*). To characterize a possible gut-sympathetic ganglia-pancreas/liver circuit, we performed polysynaptic retrograde tracing using pseudo-rabies virus (PRV). We injected GFP-expressing PRV into the pancreas and RFP-expressing PRV into the parenchyma of the liver and assessed their synaptic connections to the CG-SMG and the intestine (Fig. 4K). We detected viral spread from both organs to the CG-SMG upon dissection of intestine muscularis at day four; we observed GFP+ neurons in the myenteric plexuses of the duodenum, ileum, and colon after four days with the highest concentration of neurons in the colon and ileum, while RFP+ neurons were only observed in the ileum (Fig. 4K, fig. S7J). To investigate whether CART+ viscerofugal neuron activation could directly modulate sympathetic neuronal activity, we dissected the CG-SMG post-C21 administration and measured cFos expression as an indicator of sympathetic activation (*10, 43*). As expected, we observed a significant increase in cFos expression in C21-injected Cart^EAN-hM3Dq^ mice as compared to control animals (Fig. 4L).

To evaluate whether the above observations correlate to the microbiota-dependent changes in iEAN numbers, we quantified viscerofugal neurons in the all areas of the intestine upon microbial depletion. Indeed, retrograde fluorescent cholera toxin beta subunit (CTB) tracing from the CG-SMG revealed a preferential loss of CTB+ neurons in the ileum of antibiotic-treated mice, with no change in the colon or the sparsely retrograde–labeled duodenum (Fig. 4M, N, fig S7K,L). We measured glucose regulation in antibiotic–treated mice and found a significant reduction in blood glucose levels (Fig. 4O), results that corroborate previously reported microbiome–based modulations in glucose tolerance (*44, 45*). To determine whether microbiota–mediated changes in glucose levels are associated to loss of iEAN, we measured blood glucose in ICE^−/–^ and *Casp11*^−/–^ mice, which did not display iEAN (or CART+) loss post antibiotics treatment. In contrast to wild–type control mice, neither ICE^−/–^ nor *Casp11*^−/–^ mice showed significant changes in blood glucose levels following antibiotic treatment (Fig. 4O, fig. S7M). Finally, to directly implicate CART neurons in glucose regulation, we injected AAV5-mCherry-FLEX-DTA into the ileum and colon of *Cartpt*^Cre^ mice, to selectively delete CART+ neurons. CART+ neuron ablation resulted in a significant reduction in blood glucose and a significant increase in insulin levels as compared to *Cartpt*^Cre^ mice injected with a control AAV5 virus (Fig. 4P, Q, fig. S7N,O). These experiments demonstrate that the loss of CART+ viscerofugal iEAN can significantly impact blood glucose levels, presumably due to the lack of pancreas-specific sympathetic regulation. Together, these experiments establish a distal gut-pancreas/liver circuit that originates in microbiota-modulated CART+ viscerofugal neurons to regulate blood glucose levels.

The gut microbiota influences several physiological and pathological processes, including local nutrient absorption and lipid metabolism (*7, 44-49*), as well as activation of the gut–associated and systemic immune system (*50*). Dysbiosis or depletion of commensal bacteria has also been shown to impact iEAN excitability and neurochemical code (*51, 52*), microglia maturation (*53*), CNS neurogenesis (*54*), and behavioral or cognitive disorders (*51, 54, 55*). Our data revealed microbial– and region–dependent iEAN functional specialization with the potential to perform metabolic control independent of the CNS. Because we only focused on the functional characterization of selected neuropeptides, our study certainly does not exclude the possibility that additional microbiota-independent, modulated, or imprinted iEAN neuropeptide pathways play complementary or redundant roles in GI physiology, including in feeding behavior (*3, 45, 47, 51, 56, 57*). Nevertheless, the observation that microbiota-modulated viscerofugal neurons in the distal intestine can increase blood glucose via a local circuit warrants additional investigations into CNS-independent EAN circuits. For instance, it remains to be defined whether CART+ viscerofugal neurons respond to the presence of glucose in the lumen of the intestine, release of neuropeptides such as the incretin GLP-1, or the movement of fecal matter. Along these lines it will be important to determine whether CART+ viscerofugal neurons are functionally connected to intrinsic primary afferents, EECs, or are coupled to mechanosensation to perform glucoregulatory functions (*56, 57*). Peripheral-restricted circuits such as the one uncovered here could offer unique neuronal targets for the treatment of metabolic disorders, such as type 2 diabetes, which would bypass CNS effects.

## Methods

### Mice

Wild-type mice used: C57Bl/6J (C57Bl/6J, Jackson #000664 or C57BL/6NTac, Taconic #B6-M/F). Transgenic mice used: RiboTag (B6N.129-*Rpl22*^*tm1.1Psam*^, Jackson #011029), *Snap25*^cre^ (B6;129S-*Snap25*^*tm2.1(cre)Hze*^, Jackson #023525), *Cart*^Cre^ (B6;129S-Cartpttm1.1(cre)Hze/J, Jackson #028533), *Npy*^Cre^ (B6.Cg-Npytm1(cre)Zman/J, Jackson #027851), *Agrp*^Cre^ (Agrptm1(cre)Lowl/J, Jackson #012899), *Rosa26*lsl^-tdTomato^ (B6.Cg-*Gt(ROSA)26Sor*^*tm14(CAG-tdTomato)Hze*^, Jackson #007914). Gnotobiotic mice used: Germ-Free (GF) C57Bl/6 and *Snap25*^RiboTag^. Controls for GF C57BL6/J mice were previously GF and kept on a GF diet under SPF conditions for several generations. Controls for GF *Snap25*^RiboTag^ mice were *Snap25*^RiboTag^ SPF mice maintained on a GF diet. Mice were bred within our facility to obtain strains described and were 7-12 weeks of age for all experiments unless otherwise indicated. For comparisons to GF mice, mice were maintained on sterilized Autoclavable Mouse Breeder Diet (5021, LabDiet, USA), the same used in the gnotobiotic facility. Female mice were used for all sequencing experiments. Male and female mice were used for all other experiments. Animal care and experimentation were consistent with NIH guidelines and approved by the Institutional Animal Care and Use Committee (IACUC) at The Rockefeller University.

### Antibiotic treatments

Broad spectrum antibiotics (0.25 g Vancomycin, 0.25 g metronidazole, 0.5 g ampicillin, and 0.5 g neomycin) were dissolved in 500 mL of filtered water and supplemented with 5 g Splenda. Individual antibiotics (0.25 g vancomycin, 0.25 g metronidazole, 0.5g ampicillin and 0.5 g neomycin) were dissolved in 500 mL of filtered water and supplemented with 5 g Splenda. To control for the bitter taste of the antibiotic solution, 5 g of Splenda was dissolved in filtered water. Splenda controls were given filtered Splenda water as their drinking water. All solutions were passed through a SteriCup 0.22 um filter. Streptomycin was prepared in sterile DPBS at a concentration of 200 mg/mL and then filtered with a 0.22 uM (EMD Millipore PES Express) syringe filter. A dose of 20 mg was given as an oral gavage of 100 uL of this stock solution.

### Colonization of germ-free mice

For colonization experiments, GF mice were housed in a cage with age and sex-matched feces from SPF mice kept on a Germ Free Diet. Analyses were performed a minimum of two weeks post conventionalization.

### Virus

The following viruses were used: AAV9-hSyn-DIO-hM3Dq(Gi)-mCherry (Addgene), AAVrg-CAG-tdTomato (Addgene), AAV5-mCherry-FLEX-DTA (UNC Vector Core), AAV5-hSyn-hChR2(H134R)-EYFP (UNC Vector Core), and PRV-152/614 (Gift of L. Enquist). Fast Green (Sigma) was added (0.1%) to virus injected into peripheral tissues.

### Viral injections

Mice were anesthetized with 2% isoflurane with 1% oxygen followed by 1% isoflurane with 1% oxygen to maintain anesthesia. After shaving and sterilization of the abdomen, mice were placed on a sterile surgical pad on top of a heating pad and covered with a sterile surgical drape. Ophthalmic ointment was placed over the eyes to prevent dehydration and the incision site was sterilized. Upon loss of recoil paw compression, a midline incision was made through the abdominal wall exposing the peritoneal cavity. The duodenum, ileum, colon, or CG-SMG were located and exposed for injection. All injections were made with a pulled glass pipette using a Nanoject III. The following volumes were used for each viral injection into a different region of the intestine: AAVrg-CAG-tdTomato (1.25uL), AAV9-hSyn-DIO-hM3Dq(Gi)-mCherry (1.25uL), AAV5-mCherry-FLEX-DTA (2.5uL), and AAV5-hSyn-hChR2(H134R)-EYFP (2uL). Following injection, the abdominal wall was closed using absorbable sutures and the skin was closed using surgical staples. Antibiotic ointment was applied to the closed surgical site and mice were given 0.05 mg/kg buprenorphine every 12 h for 2 days.

### CTB viscerofugal tracing

Mice were anesthetized and operated on as described above. 1.5 DL of 1% CTB 488, 555, or 647 in PBS with 0.1% FastGreen was injected with a pulled glass pipette using a Nanoject III into the celiac-superior mesenteric ganglion. Relevant tissues were then dissected after a minimum of 2-4 days post-injection.

### Chemogenetics

Water soluble Compound 21 (HelloBio) was dissolved in sterile 0.9% saline. Mice were given an intraperitoneal injection at a dose of 1mg/kg.

### cFos counting

Mice were sacrificed by cervical dislocation and CG-SMG were harvested and fixed overnight in 4% PFA. CG-SMG were then washed four times in DPBS at RT and permeabilized in 0.5% Triton X-100/0.05% Tween-20/4 µg heparin (PTxwH) overnight RT. Primary antibody cFos (1:1000, Cell Signaling Technologies, 2250S) was added to the samples in PTxwH and incubated at 4°C for 72 h. Samples were washed four times in PTxwH at RT and then stained with goat-anti rabbit AF555/568/647 at 4°C for 48-72 h. Samples were washed four times in PTxwH at RT, covered in Fluoromount G, and coverslipped for confocal imaging. We captured all sympathetic neurons within the CG-SMG -with multiple z-stack images. All images were analyzed in Image-J. Total cFos+ nuclei were counted using the Cell Counter plugin for Image-J, and data were not normalized to area or volume. Each data point represents the number of cFos+ cells per CG-SMG.

### Brain immunofluorescence

Mice were sacrificed and transcardially perfused with cold PBS with heparin followed by cold 4% PFA (Electron Microscopy Sciences). The intact brain was separated carefully from the skull and placed in 4% PFA, and then rotated for 48 h at 4°C. Whole brains were washed with PBS/0.03%Azide and sectioned at 50 □m on a Leica vibratome for immunofluorescence. Samples were then permeabilized in 0.5% Triton/0.05 Tween-20 in PBS (PTx) followed by blocking in 5% goat serum in PTx each for 2 h at room temperature. Primary antibody was added to the blocking buffer and samples were incubated with constant rotation at 4°C overnight. Four 15-minute washes were done in PTx at RT after which samples were moved to blocking buffer with secondary antibody. Slices were incubated in secondary antibody for 2 hours at room temperature followed by four 15-minute washes in PTx at room temperature. Samples were then placed on microscope slides, covered in Fluoromount G, and coverslipped.

### Antibodies

The following primary antibodies were used, and unless otherwise indicated concentrations apply to all staining techniques: BIII-Tubulin (1:400, Millipore Sigma, T2200; 1:200, Aves Labs, TUJ), NPY (1:200, Immunostar, 22940), SST (1:400, Millipore Sigma, MAB354), RFP (1:1000, Sicgen, AB8181; 1:1000, Rockland, 600-401-379), pCREB Ser133 (1:200, Cell Signaling Technologies, 9198S), ANNA-1 (1:200,000, Gift of Dr. Vanda A. Lennon), cFos (1:1000, Cell Signaling Technologies, 2250S), HA (1:400, Cell Signaling Technologies, 3724S), CART (1:500, R&D Systems, AF163), CD9 (AF647, 1:200, BD Biosciences, 564233). Fluorophore-conjugated secondary antibodies were either H&L or Fab (Thermo Fisher Scientific) at a consistent concentration of 1:400 in the following species and colors: goat anti-rabbit (AF488/568/647), goat anti-rat (AF488/647), goat anti-chicken (AF488/568/647), goat anti-human (AF568/647), donkey anti-guinea pig (AF488/647), donkey anti-rabbit (AF568/647), donkey anti-goat (AF568/647).

### Intestine dissection

Mice were sacrificed and duodenum (4 cm moving proximal from the gastroduodenal junction), ileum (4 cm moving proximal from the ileocecal junction), or colon (4 cm moving proximal from the rectum) was removed. For AdipoClear fecal contents were flushed from the lumen and left intact. Tissue used for RIMS or FocusClear were cut open longitudinally and fecal contents were washed out. For dissection of the muscularis, following the above procedures, the intestinal tissue was placed on a chilled aluminum block with the serosa facing up(*58*). Curved forceps were then used to carefully remove the *muscularis* (*58*) in one intact sheet.

### Nodose ganglion dissection

Mice were sacrificed and the ventral neck surface was cut open. Associated muscle was removed by blunt dissection to expose the trachea and the nodose ganglion was then located by following the vagus nerve along the carotid artery to the base of the skull. Fine scissors were used to cut the vagus nerve below the nodose ganglion and superior to the jugular ganglion.

### Celiac-superior mesenteric ganglion dissection

Mice were sacrificed and a midline incision was made and the viscera were reflected out of the peritoneal cavity. The intersection of the descending aorta and left renal artery was identified, from which the superior mesenteric artery was located. The CG-SMG is wrapped around the superior mesenteric artery and associated lymphatic vessels. Fine forceps and scissors were used to remove the CG-SMG.

### Dorsal root ganglion dissection

The spinal column was isolated, cleaned of muscle, and bisected sagitally. The spinal cord was removed leaving the dorsal root ganglion (DRG) held in place by the meninges. The thoracic 13 DRG was identified by its position just caudal to thoracic vertebra. The meninges were cleared and individual DRGs were removed with fine forceps and scissors.

### RiboTag

Heterozygous or homozygous *Snap25*^RPL22HA^ were used for TRAP-seq analysis as no differences were found between either genotype. For intestine immunoprecipitation (IP) mice were sacrificed and tissue remove and divided as above. Samples were washed of fecal contents in PBS with cycloheximide (0.2 mg/mL) (PBS/CHX). Mesenteric fat was removed and the *muscularis* was separated from the mucosa as described above and samples were washed 5 times in PBS/CHX. For nodose and CG-SMG IP, tissues were isolated as described above. The RiboTag IP protocol was then followed (http://depts.washington.edu/mcklab/RiboTagIPprotocol2014.pdf) with the following modifications. All samples were homogenized by hand with a dounce homogenizer in 2.5 mL supplemented homogenization buffer (changes per 2.5 mL: 50 µL Protease Inhibitor, 75 µL heparin (100 mg/mL stock), 25 µL SUPERase• In RNase Inhibitor). Samples were then centrifuged for 10 minutes at 10,000 G, after which 800 µL of supernatant was removed and 5µL of anti-HA antibody (Abcam, ab9110) was added. Samples were kept rotating at 4°C with antibody for 1 hour. 200 µL of Thermo Protein magnetic A/G beads were washed with homogenization buffer, added to the sample, and kept rotating for 30 minutes at 4°C. The beads were washed four times with high-salt buffer and samples were eluted with 100 uL of PicoPure lysis buffer. RNA was extracted using the Arcturus PicoPure RNA isolation kit (Applied Biosystems) according to the manufacturer’s instructions.

### Ribotag RNA-sequencing

RNA libraries were prepared using SMARTer Ultra Low Input RNA (ClonTech Labs) and sequenced using 75 base-pair single end reads on a NextSeq 500 instrument (Illumina). Reads were aligned using Kallisto(*59*) to Mouse Ensembl v91. Transcript abundance files were then used in the DESeq2 R package, which was used for all downstream differential expression analysis and generation of volcano plots. For intestine samples Cre+ samples were compared with Cre-samples to generate a list of immunoprecipitated (IP) enriched genes (log2FC > 1 and padj < 0.05). This IP enriched list was then used to perform downstream analysis. Differentially expressed genes between samples were defined as those contained within the total IP enriched list from tissues being compared and with a cutoff of log2FC > 1. PCA plots were generated from log transformed DEseq2 data, as indicated in figure legends, with the FactoMineR R package. GSEA pre-ranked analysis was performed with desktop software and the C5 gene ontology database using 1000 permutations. Gene ontology enrichment analysis was performed with differentially expressed genes (log2FC > 1, padj < 0.05) using the TopGO R package and a Fisher test with an elimination algorithm was used to calculate significance.

### RNAscope whole-mount intestine immunofluorescence

C57Bl/6, *Cartp*^cre^, *Agrp*^cre^ or *Npy*^cre^ mice were sacrificed and the Duodenum, Ileum and colon removed and dissected as described above. Pieces of muscularis were pinned on sylgard coated plates and fixed in 4% PFA at room temperature for 3 hours. Samples were removed from the sylgard plates and washed in PBS twice for 10 minutes. Samples were further washed twice more in PBS or PBST for 10 minutes depending on the origin of the tissue (see table). After washing, pieces of muscularis were pinned again to sylgard plates and dehydrated along a gradient of 25/50/75/100/100 % ethanol in PBS or PBST for 10 minutes at each step (see table). 5mm x 5mm sections were cut from the tissue and mounted on slides and left to dry (∼2 minutes). Samples were digested with 50 µL of protease III digestion solution (ACDbio) at room temperature for between 30 and 45 minutes. After digestion, tissue was removed from slides using forceps and washed three times in PBS for 5 minutes each. Tissue was then hybridized using overnight at 40 °C in a humidified oven (ACDbio) using relevant probe targets. Tissue was next amplified and stained according to the RNAScope protocol for whole tissue staining with the following modifications: each amplification step was extended by 5 minutes and following the final amplification samples were washed three times for 10 minutes each. Tissue samples were mounted in Prolong gold antifade with DAPI (Thermo-Fisher). Samples were imaged within 24 hours on an inverted LSM 880 NLO laser scanning confocal and multiphoton microscope (Zeiss) and images processed using Image J.

### Whole-mount intestine immunofluorescence

Following intestine dissection, *muscularis* tissue was pinned down on a plate coated with Sylgard, followed by O/N fixation in PBS/4% PFA at 4°C. After washing in DPBS, samples were then permeabilized first in PTxwH)for 2 hours at room temperature (RT) with gentle agitation. Samples were then blocked for 2 hours in blocking buffer (PTxwH with 5% bovine serum albumin/5% donkey or goat serum) for 2 hours at RT with gentle agitation. Primary antibodies were added to blocking buffer at appropriate concentrations and incubated for 2-3 days at 4°C. After primary incubation samples were washed in PTxwH, followed by incubation with secondary antibody in PTxwH at appropriate concentrations for 2 hours at RT. Samples were again washed in PTxwH, and then mounted with FluoroMount G on slides with 1 ½ coverslips. Slides were kept in the dark at 4°C until they were imaged.

### Intestine neuronal quantification

A minimum of 10 images were randomly acquired across a piece of whole mount *muscularis*. These images were then opened in ImageJ, and the cell counter feature was used to count the number of ANNA-1+ cells in a given field. This number was then multiplied by a factor of 3.125 (25x objective), to calculate the number of counted neurons per square millimeter (mm^2^). The average of 10 (or more) images were then calculated and plotted. Thus, every point on a given graph corresponds to a single animal. For neuronal subtypes, the number of somatostatin (SST)-, CART-, neuropeptide Y (NPY)- and follistatin (FST)-positive neurons were also counted. These numbers were then reported as both number per mm^2^ and percent of ANNA-1+ neurons.

### Feeding assay

Mice were singly housed for at least 2 days prior to beginning the experiment. Before testing mice with Compound 21, feeding behavior was first assessed with saline injection during the light cycle (starting at 7:00AM) and dark cycle (starting at 19:00PM). The food intake assays were performed in the home cage. Mice were given ad libitum access to food prior to, during, and after the assay. Measurement of food intake (weighing of remaining food at each timepoint) was made at 1, 2, 4, 8, and 24 hr post i.p. injection of C21.

### Blood glucose measurement

Mice were not fasted, or fasted for either 4 or 16 hours (indicated in the figure legends) prior to analysis. Mouse tails were cut at the very tip and the first drop of blood was discarded. A single drop of blood was applied to a Breeze2 (Bayer) blood glucose test strip loaded into a Breeze2 blood glucose monitoring system (Bayer). All samples were obtained at the same time of day during the light cycle (10:00-10:30AM).

### Blood and plasma collection

Mice were not fasted, or fasted for either 4 or 16 hours (indicated in the figure legends) prior to analysis. Mouse tails were cut at the very tip and the first drop of blood was discarded. At least 100uL of blood was then collected in a Microvette (CB300) coated with Potassium/EDTA. Tubes were then centrifuged at 3600 RPM for 20 min at 4°C. Plasma was then collected and frozen at −80°C until analysis. All samples were obtained at the same time of day during the light cycle (10:00-10:30AM).

### Insulin ELISA

Insulin levels in serum samples were measured using an Ultrasensitive Mouse Insulin ELISA kit (Crystal Chem) according to the manufacturer’s instructions.

### Glucagon ELISA

Serum glucagon concentrations were determined using a Mouse Glucagon ELISA kit (Mercodia) according to the manufacturer’s protocol.

### Retrograde PRV Tracing

Mice were anesthetized and operated as described above. PRV Bartha 152 (GFP) or 614 (RFP) were a gift of L. Enquist. 3uL with 0.1% FastGreen was injected with a pulled glass pipette using a Nanoject III into the parenchyma of the right liver lobe or into the head, neck, body, and tail of the pancreas. The intestine *muscularis* and CG-SMG were harvested one to four days after injection.

### PASTAA analysis

Differentially expressed Ensembl gene ID lists (log2FC > 1, padj < 0.05) from ileum and colon samples (GF vs SPF) were used in the Predicting Associated Transcription factors from Annotated Affinities (PASTAA) web tool (http://trap.molgen.mpg.de/PASTAA.htm). Significant (p-value < 0.05) association scores for transcription factors were plotted.

### Confocal imaging

Whole mount intestine, NG, DRG, and CG-SMG samples were imaged on an inverted LSM 880 NLO laser scanning confocal and multiphoton microscope (Zeiss).

### RIMS clearing

Briefly, following secondary staining CG-SMG, nodose and DRG were submerged in Refractive Index Matching Solution (RIMS) for 30-120 min then mounted in RIMS solution on a glass slide and sealed with a coverslip for confocal imaging^29^.

### FocusClear

Whole intestine and celiac ganglion samples were first fixed in 4% PFA overnight at 4°C. Samples were then washed three times in DPBS at RT. Samples were placed into 250 µL of FocusClear solution for 15-20 minutes. They are then transferred to MountClear solution on a glass slide and a 1 ½ coverslip was used to seal the sample in place.

### AdipoClear

Adipoclear whole tissue clearing was adapted from Adipoclear protocol (*51*). Mice were sacrificed and intestinal sections were removed followed by overnight fixation in 4% PFA. Tissues were washed in PBS then dehydrated in 20/40/60/80/100% Methanol in B1N followed by dichloromethane. Tissues were then rehydrated in 100/80/60/40/20% methanol in B1N. Subsequently, samples were washed in PTxwH and then incubated in primary antibody dilutions in PTxwH for 7 Days. Samples were washed in PTxwH then incubated in secondary antibody at 1:400 in PTxwH for 7 days. Samples were again washed in PTxwH followed by PBS then dehydrated in 20/40/60/80/100% methanol followed by dichloromethane and finally cleared in dibenzyl ether.

### Light-sheet microscopy and 3D reconstruction

Whole-tissue cleared samples were imaged submerged in DBE on a LaVision Biotech Ultramicroscope II with 488 nm, 561nm or 647 nm light-sheet illumination using a 1.3x or 4x objective with 2.5um Z-slices. Images were adjusted post hoc using Imaris x64 software (version 9.1 Bitplane) and 3D reconstructions were recorded as mp4 video files. Optical slices were taken using the orthoslicer or oblique slicer tool.

### Intestine motility measurements

For measurement of total intestinal transit time, non-fasted mice were given an oral gavage of 6% carmine red dissolved in 0.5% methylcellulose (made with sterile 0.9% saline). Total intestinal transit time was measured as the time from oral gavage it took for mice to pass a fecal pellet that contained carmine. Mice in both assays were injected 2 minutes before starting with i.p. Compound 21 (1mg/kg).

### Statistical analysis

Significance levels indicated are as follows: *P < 0.05, **P < 0.01, ***P < 0.001. All data are presented as mean ± s.d.. All statistical tests used were two-tailed. The experiments were not randomized and no statistical methods were used to predetermine sample size. Multivariate data was analyzed by one-way ANOVA and Tukey’s multiple comparisons post hoc test. Comparisons between two conditions were analyzed by unpaired Student’s t-test. We used GraphPad PRISM version 8.2.0 and R 3.4.3 for generation of graphs and statistics.

## Acknowledgements

We thank all Mucida Lab members past and present for assistance in experiments, fruitful discussions and critical reading of the manuscript, Aneta Rogoz for the maintenance of gnotobiotic mice, Sara Gonzalez for maintenance of SPF mice, Tomiko Rendon and Beatriz Lopez for genotyping, the Rockefeller University Bio-imaging Research Center for assistance with the light sheet microscopy and image analysis, the Rockefeller University Genomics Center for RNA sequencing and the Rockefeller University employees for continuous assistance. We thank Jeffrey Friedman (Rockefeller University) for the generous use of lab equipment. We thank Ainsley Lockhart, Gregory Donaldson, and Veronica Jové (Rockefeller University) for critical reading of the manuscript. We also thank Michel Nussenzweig, Gabriel Victora (Rockefeller University) and Juan Lafaille (NYU) and their respective lab members for fruitful discussions and suggestions. This work was supported NIH Virus Center grant no. P40 OD010996, NIH F31 Kirchstein Fellowship (P.A.M.), NCATS NIH UL1TR001866 (P.A.M., D.M.), Philip M. Levine Fellowship (P.A.M.), Kavli Graduate Fellow (P.A.M.), Kavli Postdoctoral Fellow (M.S.), the Leona M. and Harry B. Helmsley Charitable Trust (D.M.), the Burroughs Wellcome Fund PATH Award (D.M.), Transformative R01DK116646 (D.M.).

## Author Contributions

P.A.M. initiated, designed, performed and analyzed the research, and wrote the manuscript. M.S. designed and performed experiments, analysis, and helped write the manuscript. F.M. performed experiments, analysis, figure preparation, and helped write the manuscript. Z.K. performed experiments, analysis and reviewed the manuscript. D.M. initiated, designed and supervised the research, and wrote the manuscript.

## Supplementary Video Legends

**Supplementary Video 1**. AdipoClear whole mount SPF ileum stained with anti-TUJ1.

**Supplementary Video 2.** AdipoClear whole mount GF ileum stained with anti-TUJ1.

**Supplementary Video 3.** AdipoClear whole mount SPF colon stained with anti-TUJ1.

**Supplementary Video 4.** AdipoClear whole mount GF colon stained with anti-TUJ

**Supplementary Video 5.** AdipoClear whole mount *Cartpt*^EAN-hM3Dq^ colon stained with anti-RFP.

**Fig. S1.**
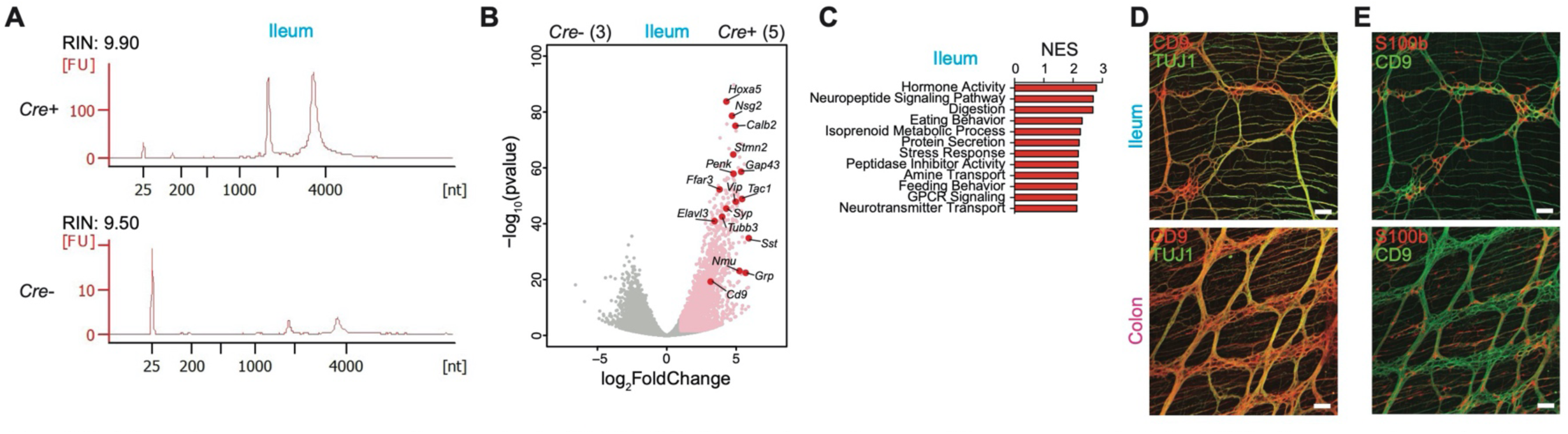
RiboTag sequencing of myenteric iEAN. (**A**) Bioanalyzer traces of immunoprecipitated (IP) ribosome-bound mRNA eluted from the ileum *muscularis* of *Snap25* ^RPL22-HA^ *Cre*+ and *Cre-* SPF mice. (**B**) Volcano plot of differentially-expressed genes from (A). Pink dots highlight those transcripts with a log2 Fold Change of >1 and padj-value <0.05. Red dots and black labels highlight selected IP-enriched genes. Number of samples shown in parentheses. (**C**) Graph of the top 12 gene ontology pathways with a FDR < 25% identified by Gene Set Enrichment Analysis (GSEA) from ileum IP-enriched genes (log2 Fold Change > 1, padj < 0.05). NES, normalized enrichment score; FDR, false detection rate. (**D, E**) Representative whole-mount immunofluorescence image of the ileum and colon myenteric plexus of C57BU6J SPF mice stained with anti-CD9 (red) and anti-neuronal-specific TUJ1 (green) (D), or anti-CD9 (green) and anti-glia-specific S100b (red) (E). Scale bars= 50 µm. Images representative of n = 3.

**Fig. S2.**
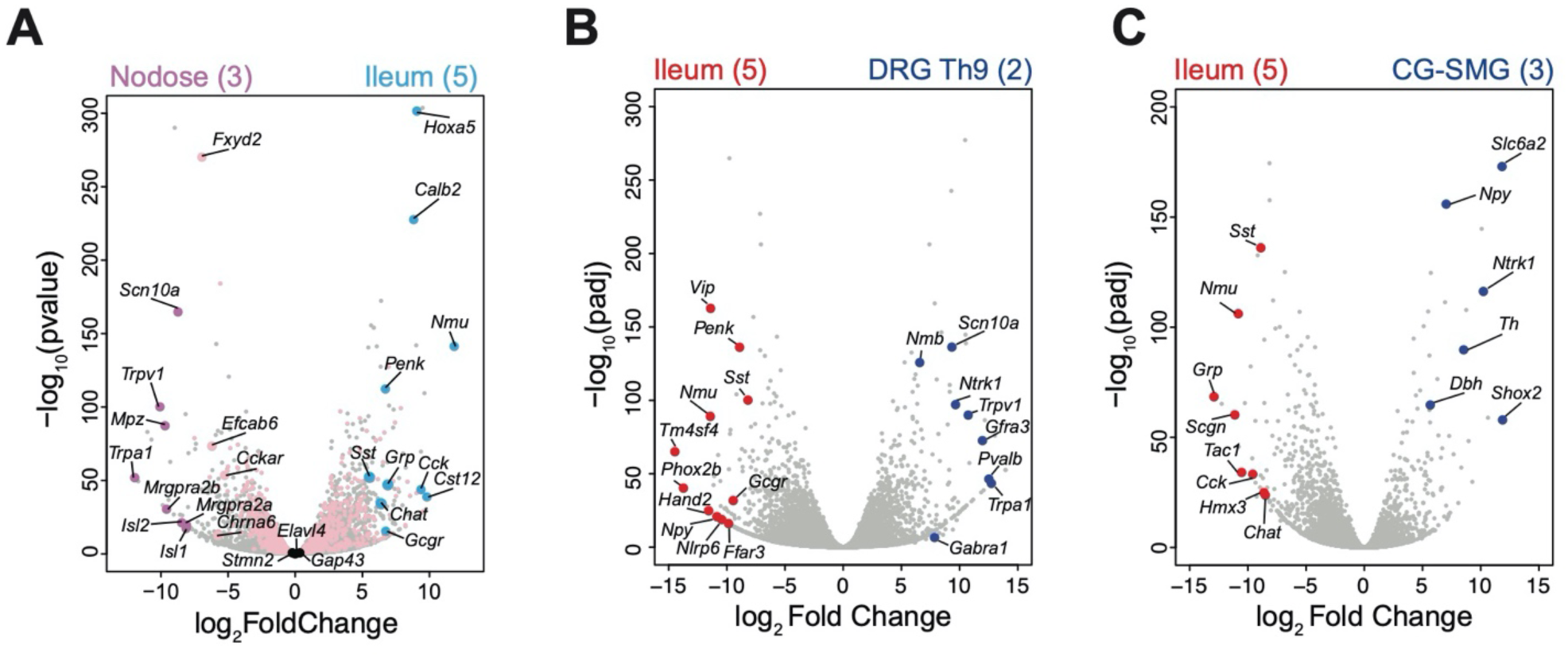
TRAPseq comparison of iEAN and eEAN. (**A**) Volcano plot of differential ly-expressed genes from the nodose ganglion (A), DRG Th9 (B), CG-SMG (C) and ileum mynteric iEAN of Snap25^RiboTag^ SPF mice. Grey dots highlight all genes analyzed by differential expression analysis. In (A) pink dots highlight all IP-enriched genes from the ileum. Purple and blue dots highlight differentially expressed genes from nodose gangli on and ileum, respectively. Enlarged pink dots represent ileum IP-enriched transcripts expressed at higher levels than nodose. Black dots highlight neuronal genes that do not significantly differ between samples. In (B), enlarged red dots highlight genes enriched in the ileum. Enlarged blue dots highlight genes enriched in DRG. In (C), enlarged red dots highlight genes enriched in the ileum. Enlarged blue dots highlight genes enriched in CG-SMG. Sample numbers are indicated in parentheses at top.

**Fig. S3.**
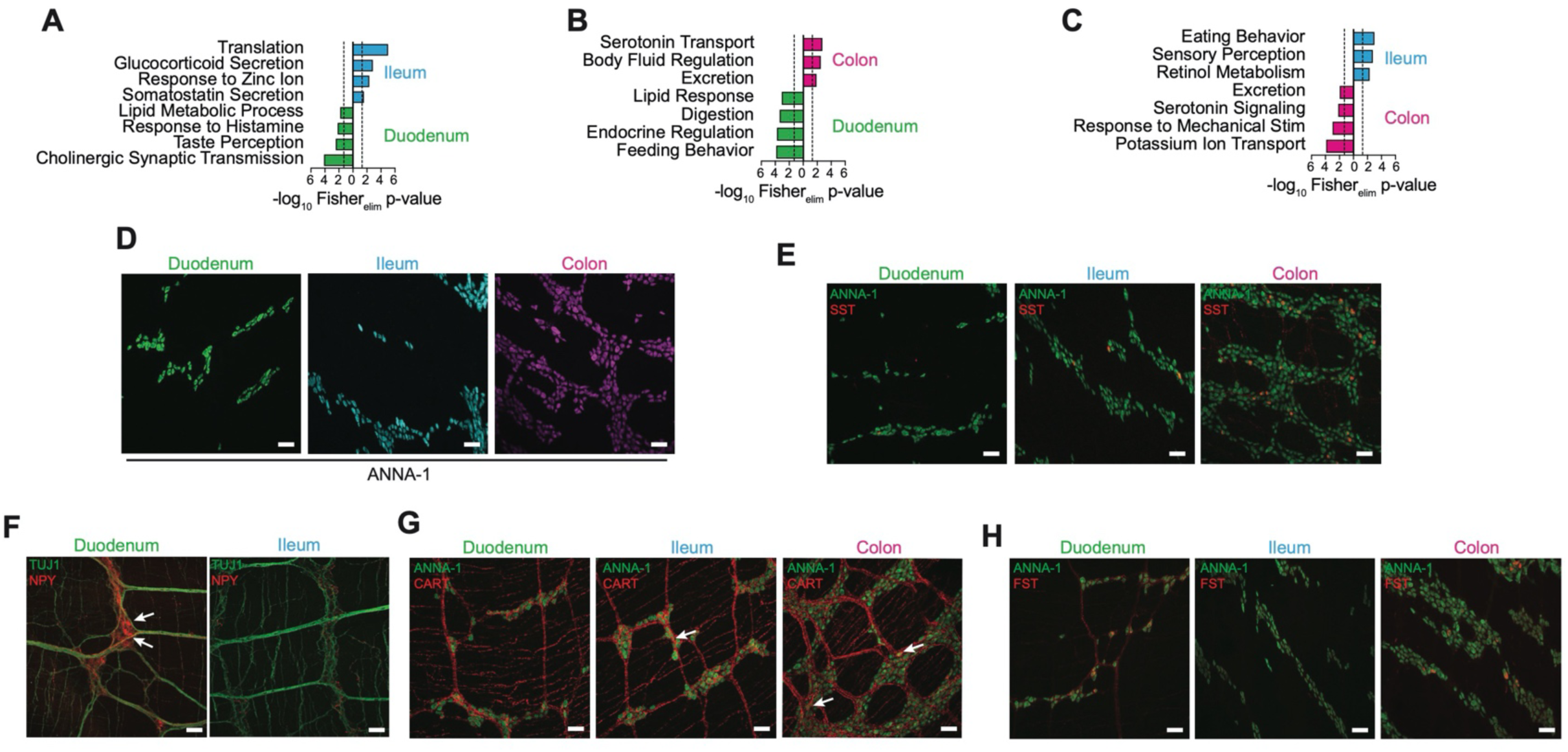
iEAN anatomical differences in gene pathways and specific genes. (**A**) Gene ontology pathways, identified by TopGO analysis of differentially-expressed genes (log2 Fold Change > 1, padj < 0.05), enriched in the ileum (blue) vs duodenum (green). Dashed lines represent threshold of significance (1.3) as calculated by Fisher’s test with an elimination algorithm. (**B**) Gene ontology pathways, identified by TopGO analysis of differentially-expressed genes (log2 Fold Change > 1, padj < 0.05), enriched in the colon (magenta) vs duodenum (green). Dashed lines represent threshold of significance (1.3) as calculated by Fisher’s test with an elimination algorithm. (**C**) Gene ontology pathways, identified by TopGO analysis of differentially-expressed genes (log2 Fold Change > 1, padj < 0.05), enriched in the ileum (blue) vs colon (magneta). Dashed lines represent threshold of significance (1.3) as calculated by Fisher’s test with an elimination algorithm. (**D**) Representative whole-mount immunofluorescence (IF) images of myenteric iEAN (stained with anti-ANNA-1) in the duodenum, ileum, and colon. Scale bar = 50 µm. (**E**) Representative whole-mount IF images of SST+ (red) myenteric iEAN in the duodenum, ileum, and colon. Scale bar = 50µm. (**F**) Representative whole-mount IF images of neuropeptide Y (NPY)+ (red) myenteric iEAN in the duodenum and ileum. Scale bar = 50µm. (**G**) Representative whole-mount IF images of CART+ (red) myenteric iEAN in the duodenum, ileum, and colon. Scale bar = 50 µm. (**H**) Representative whole-mount IF images of follistatin (FST)+ (red) myenteric iEAN in the duodenum, ileum, and colon. Scale bar = 50 µm.

**Fig. S4.**
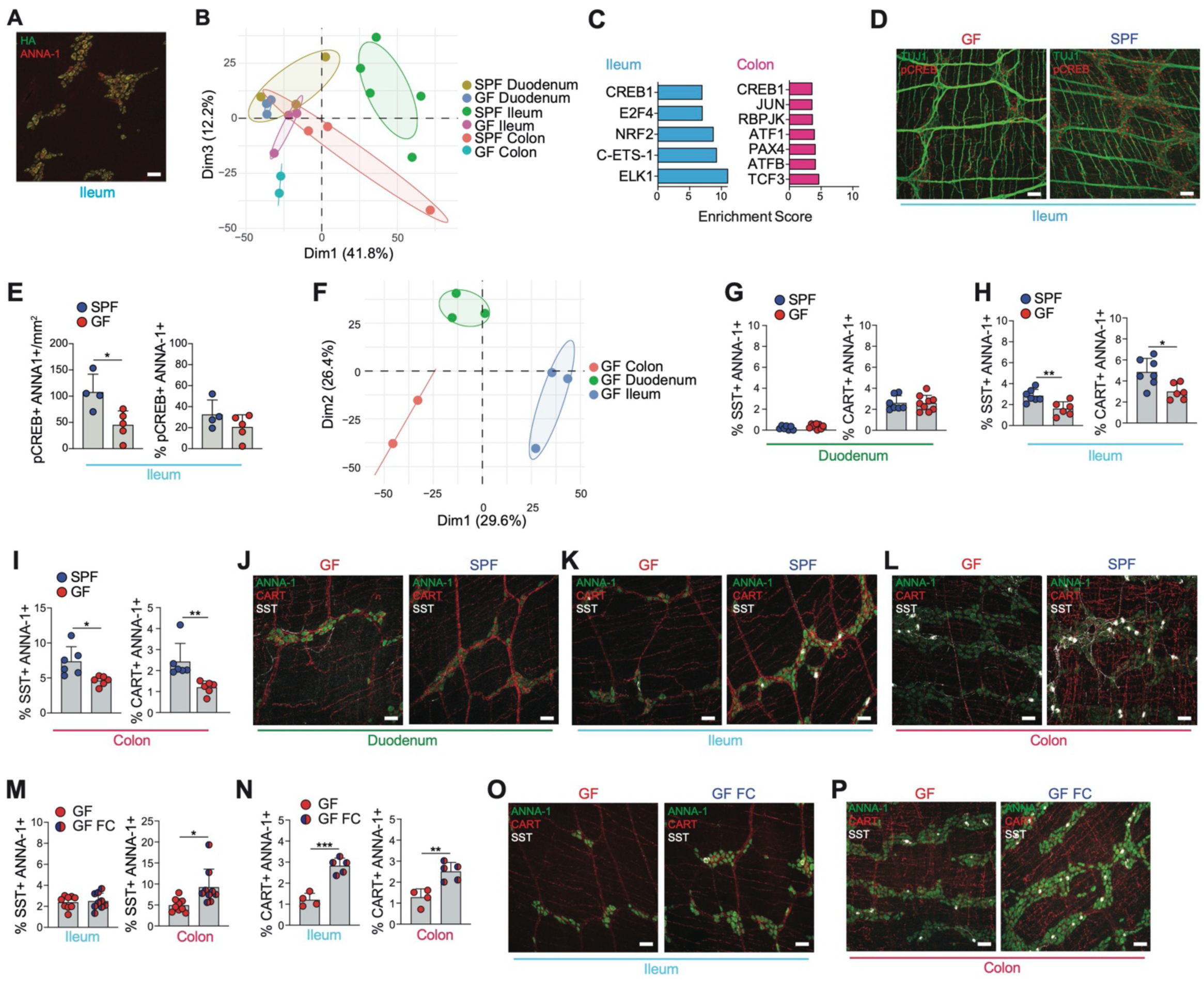
Microbiota impacts iEAN translatome in a compartimentalized manner. (**A**) Expression of HA in ileum myenteric neurons of *Snap25*^RPL22-HA^ GF mice. Scale bar = 50 µm. Image representative of duodenum and colon mytenteric plexus. (**B**) Principal Component Analysis of GF and SPF iEAN from the duodenum, ileum, and colon. Total comparative gene list generated from IP enriched transcripts (log2Fold Change > 2.5, padj < 0.05) from each sample. (**C**) PASTManalysis showing enrichment of predicted CREB1 transcription factor control of the most singificant differentially-expressed genes (log2Fold Change > 1, padj < 0.05) of the ileum and colon iEAN between C57BL/6J GF and SPF mice. (**D**) lmmunofluorescent (IF) staining of the ileum myenteric plexus from C57BU6J GF (left) and SPF (right) mice using anti-pCREB (red) and anti-TUJ1 (green) antibodies. Scale bars = 50 µm. (**E**) Number and percentage of pCREB+ myenteric iEAN in the ileum of GF and SPF mice. * *P* < 0.05 as calculated by unpaired t-test. (**F**) Principal Component Analysis of GF iEAN from the duodenum, ileum, and colon. Lmmuno-precipitated (IP) transcripts (log2 Fold Change > 1, padj < 0.05) were used to generate the list of genes for comparison between all groups. (**G-1**) Percentage of SST+ (left) and CART+ (right) myenteric iEAN in the duodenum (G), ileum (H), and colon (I) of GF and SPF mice. * *P* < 0.05, ** *P* < 0.01 as calculated by unpaired t-test. (**J-L**) Whole mount IF images of somatostatin (SST)+ and CART+ myenteric iEAN in the duodenum (J), ileum (K), and colon (L) of GF (left) and SPF (right) mice. Scale bar = 50µm. (**M** and **N**) Percentage of somatostatin (SST)+ (M) and CART+ (N) myenteric iEAN in the ileum (left), and colon (right) of GF and SPF mice. * *P* < 0.05, ** *P* < 0.01, *** *P* < 0.001 as calculated by unpaired t-test. (**O** and **P**) Representative whole-mount IF images of SST+ and CART+ myenteric iEAN in the ileum (O) and colon (P) of GF (left) and GF mice colonized with SPF microbiota (fecal colonozation, GF FC, right). Scale bar = 50µm.

**Fig. S5.**
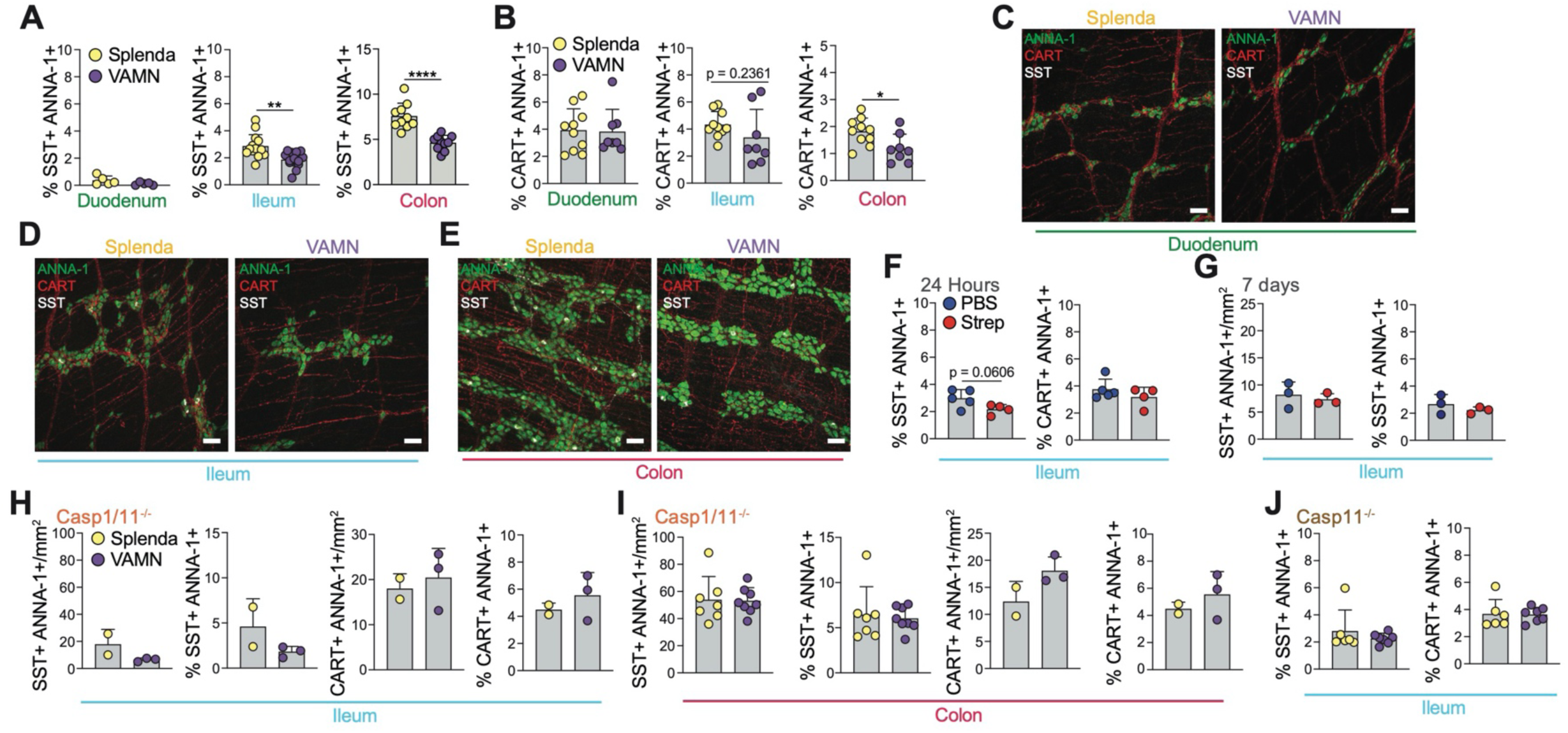
Microbial depletion leads to iEAN neuropeptide changes. (**A** and **B**) Percentage of somatostatin (SST)+ (A) and CART+ (B) myenteric iEAN in the duodenum, ileum, and colon of C57BL6/J mice treated with broad-spectrum antibiotics (vancomycin, ampicillin, metronidazole and neomycin - VAMN) administered in Splenda-supplemented drinking water or Splenda (artificial sweetener) only in the drinking water for two weeks.* *P* < 0.05, ** *P* < 0.01, **** *P* < 0.0001 calculated by unpaired t-test. (**C-E**) lmmunofluorescence whole-mount images of myenteric iEAN stained with ANNA-1 (green), CART (red), and somatostatin (SST) (white) of mice treated with Splenda (left) or broad-spectrum antibiotics (VAMN, right) for 2 weeks, in the duodenum (C), ileum (D), and colon (E). Scale bars= 50 µm. Images representative of at least n=5/group. (**F** and **G**) Percentage and number of SST+ and CART+ myenteric iEAN in the ileum of C57BL6/J mice treated with one oral gavage of streptomycin and analyzed 24 hours (F) or 7 days (G) post gavage. (**H** and **I**) Number and percentage of somatostatin (SST)+ (left) and CART+ (right) myenteric iEAN in the ileum (H) and colon (I) of *Casp1/11*^*-/-*^*mice* treated with Splenda or VAMN drinking water for 2 weeks. (**J**) Percentage of SST+ (left) and CART+ (right) myenteric iEAN in the ileum *Casp11*^*-/-*^*mice* treated with Splenda or broad-spectrum antibiotics (VAMN) in the drinking water for 2 weeks.

**Fig. S6.**
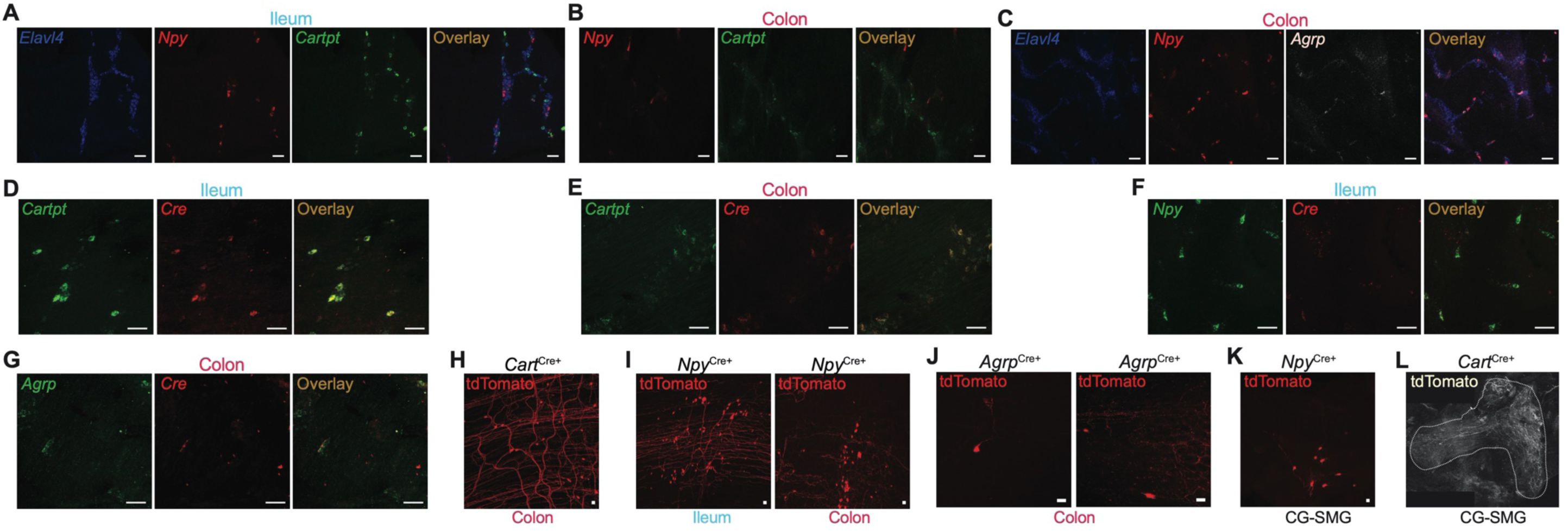
Characterization of neuropeptide expression and patterning in the distal intestine. (**A-C**) RNAscope *in situ* hybridization immunofluorescence whole-mount images of the indicated intestinal segments from C57BL6/J mice using probes for (A-B) *Elavl4, Npy*, and *Cartpt* in the ileum (A) and colon (B) or *Elavl4, Npy*, and *Agrp* in the colon (C). Scale bars = 50 µm. (**D-G**) RNAscope *in situ* hybridization immunofluorescence whole-mount images of intestinal segments from *Cart*^Cre^ (D-E), *Npy*^Cre^ (F), and *Agrp*^Cre^ (G) of the ileum (D,F) and colon (E,G) using probes for *Cre* and (D-E) *Cartp*, (F) *Npy* and (G) *Agrp.* Scale bars = 50 µm. (**H-I**) lmmunofluorescence whole-mount images of the (H) the colon from *Cart*^Cre^ mice, and (I) the ileum (left) and colon (right) of *Npy*^Cre^ mice injected with AAVrg-FLEX-tdTomato into the duodenum, ileum, and colon. Scale bar = 50 µm. (**J**) lmmunofluorescence whole-mount images of the colon from *Agrp*^Cre^ mice injected with AAVrg-FLEX-tdTomato into the mid-colon. Scale bars = 50 µm. (**K-L**) lmmunofluorescence whole-mount image of the CG-SMG from (K) *Npy*^Cre^ and (L) *Cart*^Cre^ mice injected with AAVrg-FLEX-tdTomato into the duodenum, ileum, and colon. Scale bars = 50 µm.

**Fig. S7.**
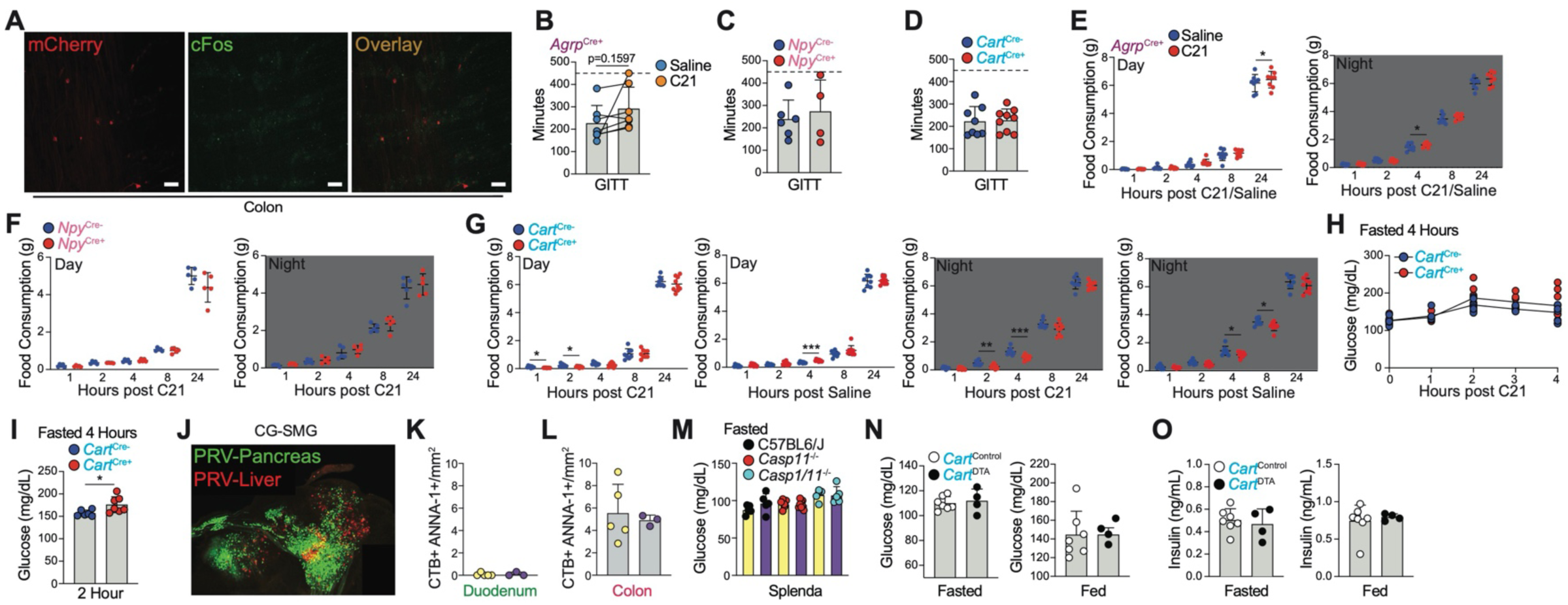
Comparison of distal intestine neuropeptide iEAN function. (**A**) Whole-mount immunofluorescence (IF) image 3 hours post administration of Compound 21 (C21, 1mg/kg) of cFos (green, stained with antibody) and mCherry (red, stained with antibody) in the colon myenteric plexus from *Agrp*^Cre^ mice injected with AAV9-hSyn-DIO-hM3Dq-mCherry into the mid-colon. Scale bars = 50 µm. (**B-D**) Gastrointestinal transit time (GITT) of *Agrp*^Cre^ (B), *Npy*^Cre^ (C), and *Cart*^Cre^ (D) mice injected with AAV9-hSyn-DIO-hM3Dq-mCherry into the ileum and colon. Non-significant p-value as calculated by paired I-test (B). (**E-G**) Food consumption during the day (left) or at night (right) post administration of compound 21 (C21, 1mg/kg) or saline in *Agrp*^Cre^ (E), *Npy*^Cre^ (F), *Cart*^Cre^ (G) mice injected with AAV9-hSyn-DIO-hM3Dq-mCherry into the ileum and colon. * *P* < 0.05, ** *P* < 0.01, *** *P* < 0.001 as calculated by unpaired t-test. (**H**) Blood glucose levels of *Cart*^Cre^ mice fasted for 4 hours followed by administration of C21 (1mg/kg), injected with AAV9-hSyn-DIO-hM3Dq-mCherry into the ileum and colon 2 weeks prior to analysis. (**I**) Blood glucose levels of *Cart*^Cre^ mice fasted for 4 hours, followed by administration of C21 (1mg/kg), and analyzed 2 hours later. Mice were injected with AAV9-hSyn-DIO-hM3Dq-mCherry into the ileum and colon 2 weeks prior to analysis. * *P* < 0.05 as calculated by unpaired t-test. (**J**) Whole-mount IF image of the CG-SMG 4 days post injection of PRV-GFP into the pancreas and PRV-RFP into the liver. (**K-L**) Number of CTB+ neurons in the duodenum (K) and colon (L) after injection of CTB-AF647 into the CG-SMG of C57BU6J mice treated with broad-spectrum antibiotics (vancomycin, ampicillin, metronidazole and neomycin - VAMN) administered in Splenda-supplemented drinking water or Splenda (artificial sweetener, control) in the drinking water for two weeks. (**M**) Blood glucose levels of C57BL6J, *Casp1/11*^−*/*−^, and *Casp11*^−*/*−^ mice treated with Splenda (artificial sweetener, control) in the drinking water before switching experimental groups on drinking water containing broad-spectrum antibiotics (VAMN). (**N**) Blood glucose levels of fasted (left) or fed (right) *Cart*^Cre^ mice before injection with AAV5-mCherry-FLEX-DTA or control virus. (**O**) Plasma insulin levels of fasted (left) or fed (right) *Cart*^Cre^ mice before injection with AAV5-mCherry-FLEX-DTA or control virus.

